# The Arabidopsis TTL3 protein interconnects brassinosteroid signalling and cytoskeleton during lateral root development

**DOI:** 10.1101/2020.12.21.423430

**Authors:** Pengfei Xin, Jakub Schier, Ivan Kulich, Joseph G. Dubrovsky, Vielle-Calzada Jean-Philippe, Aleš Soukup

## Abstract

Lateral roots are essential components of the plant edaphic interface, contributing to water and nutrient uptake, biotic and abiotic interactions, stress survival, and plant anchorage. We have identified the TETRATRICOPEPTIDE-REPEAT THIOREDOXIN-LIKE 3 (TTL3) being related to lateral root emergence and later development. TTL3 interacts with microtubules and potentially interconnects cytoskeletal function with the brassinosteroid signalling pathway. Loss of function of *TTL3* leads to a reduced number of emerged lateral roots due to delayed development of lateral root primordia. Lateral root growth of the *ttl3* mutant is less sensitive to BR treatment. Timing and spatial distribution of *TTL3* expression is consistent with its role in development of lateral root primordia before their emergence and subsequent development into lateral roots. TTL3 is a novel component of the root system morphogenesis regulatory network.

## Introduction

Root system architecture determines plant efficacy during uptake of water and inorganic nutrients, stress survival, and biotic interactions. In seed plants, most of the root system absorptive surface is formed by lateral roots, originating mostly post embryonically from the pericycle of the parent root of seed plants (Malamy and Benfey, 1997; Hochholdinger and Zimmermann, 2008). Root branching increases plant contact with the soil environment, which has implications for plant performance under various environmental conditions such as drought, salinity, nutrient deficiency (Vilches-Barro and Maizel, 2015; Stoeckle et al., 2018; von Wangenheim et al., 2020).

A complex developmental network coordinates root branching – maximizing plants’ ability to adapt to ever changing environmental conditions. There is a wide set of genes of the regulatory circuit oscillating during the lateral root initiation (Delay et al., 2013). The auxin signalling takes a central role in lateral root primordium (LRP) initiation and development (Fukaki et al., 2002; Wilmoth et al., 2005; Du and Scheres, 2018; Xuan et al., 2020). Lateral root development is further influenced by mechanical signalling between the lateral root and adjacent maternal tissues (Stoeckle et al., 2018). Endodermal cells accommodate the volume changes in the developing primordium (Marhavý et al., 2016). Spatial distribution, timing of cell division, and radial asymmetric expansion of cells (all of which vary according to position within a LRP) determine the shape of the LRP (Stoeckle et al., 2018). Volume growth of individual cells relies on tissue context, biomechanical properties of cell walls, and the geometry of the cell. During the emergence of the LRP from its maternal tissue, penetration through overlaying tissues is facilitated by cell wall modifying enzymes (Swarup et al. 2008). Cell wall properties are largely determined by the orientation of cellulose microfibrils (Lloyd et al., 1985; Baskin et al., 2004; Landrein and Hamant, 2013). Pectin esterification and related cell wall mechanical properties are essential for proper LRP morphogenesis (Wachsman et al., 2020) and most likely, lateral root emergence. In both cases, cortical cytoskeleton is thought to be strongly involved. Cortical microtubules direct cellulose synthase complexes to deposit cellulose microfibrils (Paredez et al, 2006; Gutierrez et al, 2009; Landrein and Hamant, 2013). They also target the vesicle transport of non-cellulosic cell wall components and pectin modifying enzymes (Kim and Brandizzi, 2016). There is an intriguing link between pectin metabolism and brassinosteroid signal transduction (Wolf et al., 2014, Wachsman et al., 2020).

Brassinosteroids (BRs) regulate various aspects of lateral root development via a complex regulatory network. The effect of BRs is thought to occur through interactions with auxin (Kim et al., 2000; Bao et al., 2004; Li et al., 2005; Kim et al., 2007; Chaiwanon and Wang, 2015). BRs modulate auxin’s acropetal transport and distribution within the roots (Bao et al., 2004; Li et al., 2005). Moreover, auxin induces the expression of the brassinosteroid biosynthetic gene DWF4 in roots; in *dwf4* mutants, auxin-induced lateral root elongation was suppressed (He et al., 2005; Yoshimitsu et al., 2011). BRASSINOSTEROID INSENSITIVE 2 (BIN2) regulates lateral root organogenesis by phosphorylating ARF7 and ARF19 (Cho et al., 2014; Maharjan et al., 2011). BIN2 activity is inhibited by BRs via activation of the BZR1 and BES1 transcription factors (He et al., 2002; Wang et al., 2002; Yin et al., 2002). Exogenous application of epibrassinolide (eBL) induces expression of several Aux/IAA genes involved in root development (Kim et al., 2006). The auxin-insensitive *iaa7/axr2-1* and *iaa17/axr3-3* mutants show increased eBL sensitivity in roots (Nakamura et al., 2006).

The role of TTL proteins in lateral root development is not yet understood. The TTL gene family is specific for terrestrial plants and is anticipated to play a role in plant development and environmental interactions (Rosado et al., 2006; Lakhssassi et al., 2012; Amorim-Silva et al., 2019).

Here, we report further functional characterization of the TTL3 (TETRATRICOPEPTIDE-REPEAT THIOREDOXIN-LIKE 3) gene. Cytoplasmic TTL3 protein has been proven to interact with BR signaling components and modulate growth response in the presence of osmotic stress and salinity (Lakhssassi et al, 2012; Amorim-Silva et al., 2019). To further understand its effects in roots, we have characterized knockout mutant *ttl3*, which has delayed LRP development and a reduced number of emerged lateral roots. After BR treatment, the mutant *ttl3* was less affected than the wild type. Our data prove that TTL3 protein interacts with microtubules and is likely interconnected with vesicle transport in Arabidopsis cells. The involvement of TTL3 in lateral root development interconnects the cytoskeleton and BR signalling.

## Materials and Methods

### Plant Materials and Growth Conditions

*Arabidopsis thaliana*, Columbia ecotypes (Col-0) were used as a corresponding WT to insertion knock-out mutants *ttl1* (SALK 063943), *ttl3* (SAIL193B05), and *ttl4* (SALK 134098), which were obtained from Nottingham Arabidopsis Stock Centre (NASC). The homozygous lines *TTL3* (wild type) and *ttl3* were selected after the heterozygous SAIL193B05 self-cross. Plant genotypes were identified by PCR using standard primers from signal.salk.edu. Mutants *ttl1* and *ttl3, ttl1* and *ttl4*, and *ttl3* and *ttl4*, were crossed to obtain the double mutants *ttl1ttl3, ttl1ttl4*, and *ttl3ttl4*. Double mutants *ttl1ttl3* and *ttl1ttl4*, and *ttl1ttl3* and *ttl3ttl4* were hybridized to obtain the triple mutants *ttl1ttl3ttl4-1* and *ttl1ttl3ttl4-2*, respectively. Complemented line cTTL3 was obtained after transformation of *pTTL3::TTL3-GFP* into *ttl3* plants. The Arabidopsis seeds were surface-sterilized and vernalized in sterile water in the dark at 4 °C for 3 days. Plants were grown on sterile agar Petri dishes inclined at 45°, containing 0.2× MS salts (Murashige and Skoog, 1962), 1% (w/v) sucrose, and 1% (w/v) agar. These were grown at 22 °C under long-day conditions (8h-dark/16h-light cycle) in a cultivation room. For Epibrassinolide (eBL) treatment, seedlings were germinated as described above and transferred after 5 days to sterile agar supplemented with 100 nM eBL. Petri dishes were scanned 6 days later. For the LRP development analysis, 1/2 MS plates without sucrose were used (Deng et al., 2017).

### Constructs and Plant Transformation

The TTL3 promoter region, 2.5 kb upstream of the start codon, was amplified (primers pTTL3-LP and pTTL3-BP, Table S1) and subcloned into the pENTR5’-TOPO TA Cloning vector (Invitrogen). The gene sequence of *TTL3* gDNA was amplified (primers TTL3-gDNA-LP and TTL3-gDNA-BP, Table S1) and subcloned into the pDONR221 (Invitrogen). GFP CDS was cloned into the pEN-R2-F-L3.0 vector (Karimi et al., 2007). Both were combined into the Gateway binary vector pK7m34GW to gain the *pTTL3::TTL3-GFP* construct. To generate *pTTL3::GUS* transcriptional fusion, the *TTL3* promoter was introduced into the *pMDC162* vector. The plasmids containing *pTTL3::TTL3-GFP* and *pTTL3::GUS* were transfected into Col-0 or *rdr6* (Scholl et al., 2000) background of Arabidopsis using *Agrobacterium tumefaciens* GV3101 by floral dip (Clough and Bent, 1998). Hygromycin (30μg/mL) or BASTA (150mg/L) herbicide treatment were used to select transgenic plants. Transient transformation of 5 to 8 weeks old plants of *Nicotiniana benthamiana* was carried out by *Agrobacterium tumefaciens* containing *35S::TTL3-GFP* plasmid and viral p19 post-translational silencing suppressor (Qiu et al. 2002). To introduce microtubule marker into *pTTL3::TTL3-GFP* line, the plants were additionally transformed with *35S::MAP4-RFP* (Microtubule-associated protein 4) (Marc et al., 1998), both in Col-0 and *rdr6* background. The *rdr6* Arabidopsis mutant is deficient in RNA-dependent RNA Polymerase 6 (RDR6), which takes part in posttranscriptional gene silencing. The *rdr6* genetic background is therefore suitable for the expression of potentially silenced genes (Butaye et al., 2004; Sasse et al., 2015). The tobacco (*N. benthamiana* containing *35S::TTL3-GFP*) and Arabidopsis (containing *pTTL3::TTL3-GFP*) were treated with 1mM and 100μM oryzalin, respectively, to follow changes induced by depolymerization of microtubules.

**Table S1.**
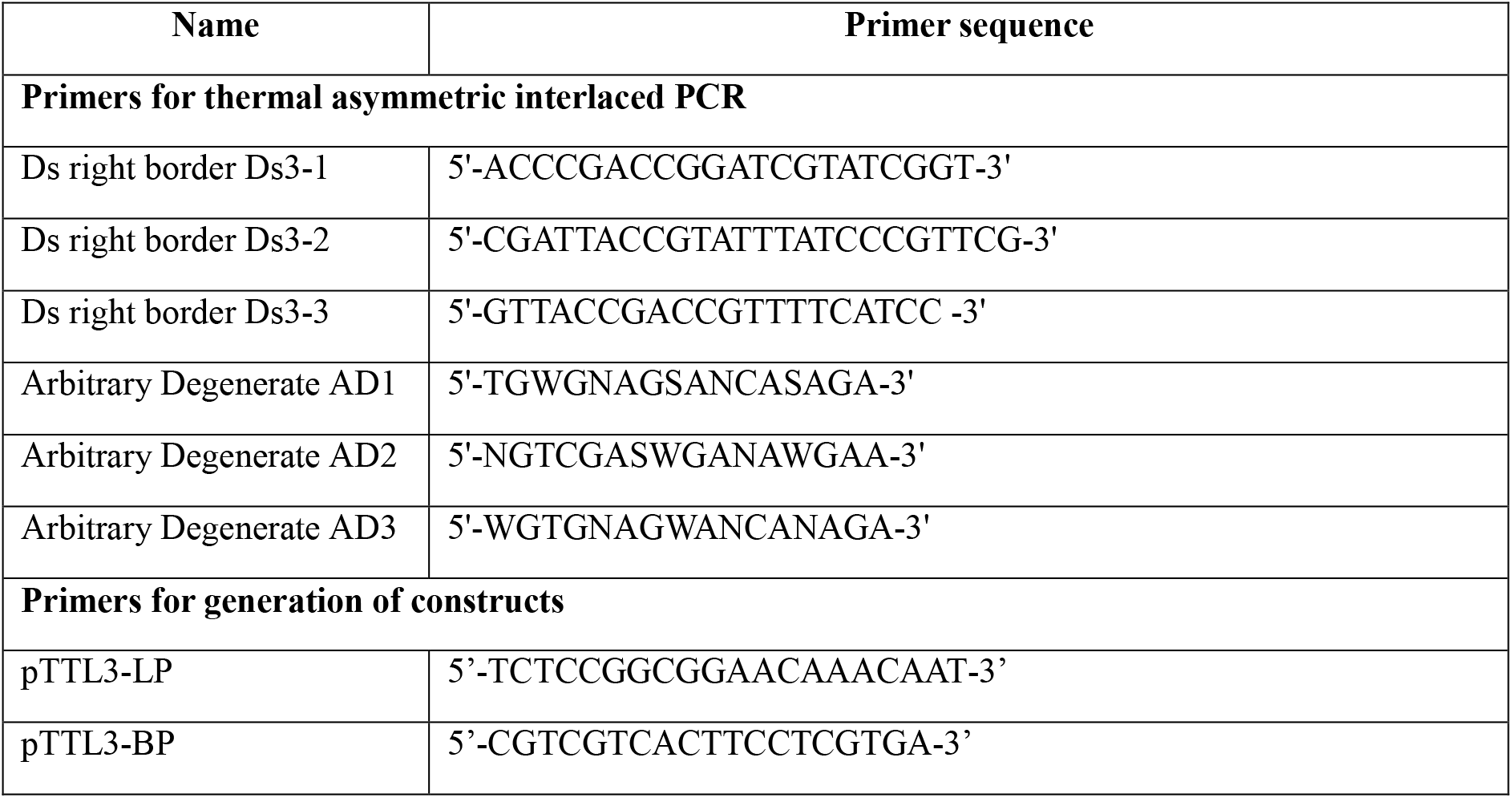
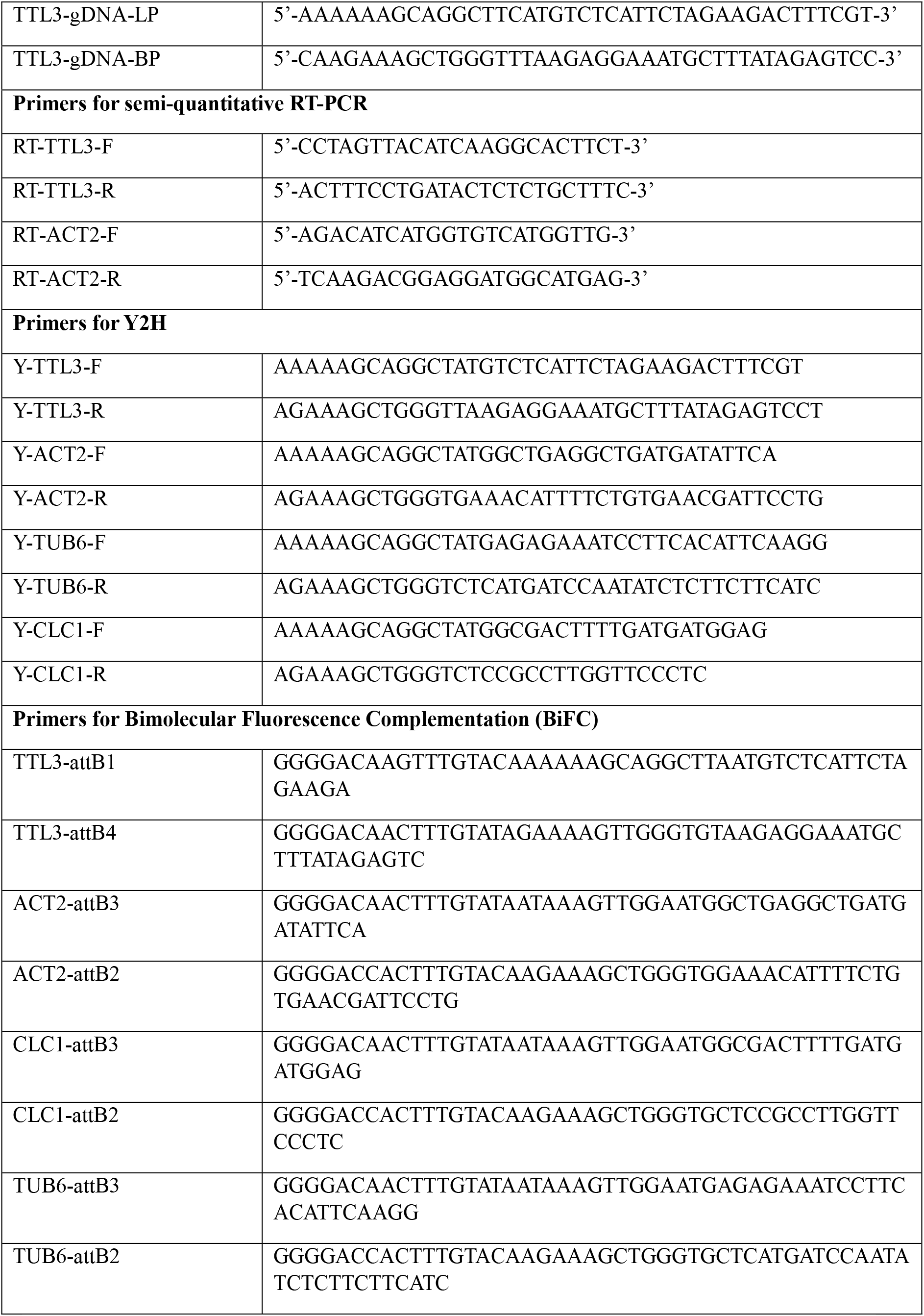
List of primers

### RNA Isolation and semiquantitative RT-PCR Analyses

Total RNA was extracted from 10-day-old Col-0, *TTL3*, and *ttl3* seedlings, using an RNeasy RNA plant extraction mini kit (Qiagen, GmbH, GERMANY) according to the manufacturer’s protocol. To synthesize the first strand of cDNA, 2 μg of total RNA was used for reverse transcription. The Arabidopsis actin 2 gene (AT3G18780) was used as the internal standard. For primers used see Table S1.

### Root morphological analysis and LRP Development

Root systems were scanned 7, 8 and 9 days after germination at 1200 DPI. Quantification of the total root system length, primary root length, total lateral roots length, and number of lateral roots was performed using the Root Analyzer module in NIS-elements software (LIM Prague, CZ). The lateral root density was calculated from the number of lateral roots and the branched zone length. The root apical meristem (RAM) size was estimated by measuring the mean value of the number and length of cortical cells between the quiescent center (QC) and the first cell in the elongation zone (Pavelescu et al., 2018). Phenotypic trait analysis was performed from at least two independent experiments with a minimum of 10 replicates each.

To observe the rate of development of LRP, 5-day-old seedlings grown on vertical 1/2 MS plates without sucrose were rotated by 90° to stimulate the simultaneous production of LRP (Lucas et al., 2008). The seedlings underwent gravitropic stimulation for 20h or 50h and the induced LRP was microscopically scored under DIC optics (Olympus BX 51). For statistical analysis of the developmental stage distribution of LRPs, the Pearson’s independence test was used to compare pairs of LRP developmental stages in the relevant genetic background (Zar, 2010). In the eBL-treated root bending assay, seedlings were grown on vertical 1/2 MS plates without sucrose for 5 days, then transferred to sucrose-free MS medium without and with 100nM eBL respectively. After one-day cultivation, the seedlings were rotated by 90° to stimulate synchronised production of LRPs. Those were evaluated 50h after gravitropic stimulation.

### The 5-ethynyl-2-deoxyuridine (EdU) cell proliferation assay

An EdU-based assay was performed to detect the number of cells in the S-phase of the Arabidopsis primary root tip (Kotogány et al., 2010), using EdU-Click 488 (Baseclick GmbH, 82061 Neuried, Germany). The 7-day-old seedlings were treated with 20μM EdU for 45 minutes, fixed in 4% (w/v) formaldehyde and labelled according to manufacturer protocol. Number of labelled nuclei were estimated from 20 optical sections (Z distance −1μm) using 3-D thresholding (3-D ImageJ suite - https://imagej.net/3D_ImageJ_Suite).

### Microscopic Analysis

A GUS assay was performed on plants fixed in the 90% Acetone and cooled to −20°C. Stained plants were mounted in clearing media (Soukup, 2014) for observation.

OLYMPUS BX51 and Apogee U4000 camera, Nikon ECLIPSE 90i fit with Andor Zyla 5.5 w and Zeiss LSM880 confocal microscope and CSU-X1 Yokogawa spinning disk head and Photometrics Prime 95B camera fitted to a Nikon Ti-E inverted microscope were used for imaging. NIS-elements, Zeiss ZEN, and ImageJ (https://imagej.nih.gov/ij/) programs were used for image acquisition and processing. The image processing Images were acquired in 16bit depth and adjustments (brightness, Gamma 0,8) were applied to the entire image (Fig. 6 and 7).

### Co-immunoprecipitation and Mass Spectrometry Analysis

Total proteins were extracted from 2-week-old Arabidopsis seedlings expressing *pTTL3::TTL3-GFP* and *35S-GFP* with 20 mM HEPES buffer (pH 6.8, 150 mm NaCl, 1 mm EDTA, 1 mm DTT, and 0.5% Tween), supplemented with a protease inhibitor cocktail (Sigma-Aldrich). The protein extracts were centrifuged at 10,000 g for 10 min at 4 °C. Protein complexes with TTL3-GFP and free GFP as a control were isolated using the μMACS and MultiMACS GFP Isolation Kits (Miltenyi Biotec, Germany), according to the manufacturer’s instructions. The anti-GFP microbeads were added and incubated for 30 min on ice with occasional shaking. The sample on the column was washed by lysis buffer and immunoprecipitates were eluted with 50μL of the elution buffer. Western blot was used to confirm the presence of fusion protein.

Mass spectrometry was carried out using UHPLC Dionex Ultimate3000 RSLC nano (Dionex, Germany) connected with the mass spectrometer ESI-Q-TOF Maxis Impact (Bruker, Germany). For protein identification, Mascot server (version 2.4.1, Matrix Science) was used. Mascot decoy search was used to calculate false discovery rate (FDR). Identified proteins were filtered so that the final FDR was less than 1%.

### Yeast Two-Hybrid Assay

The open reading frame of *TTL3* was fused to the pGADT7 vector. Preselected candidates for interacting proteins (TUB6-AT5G12250, ACT2-AT3G18780, CLC1-AT2G20760) were cloned into the pGBKT7 vector using the primers shown in Table S1. The vectors were transformed into the Y2H Gold Yeast Strain. Briefly, yeast transformed with the respective constructs were selected first on plates without leucine and tryptophan and then in a 10× dilution series starting with OD600 nm=0.1 on plates without leucine, tryptophan, and adenine to test for protein-protein interactions (Brückner et al., 2009).

### Bimolecular Fluorescence Complementation Assay

For the bimolecular fluorescence complementation (BiFC) assay (Kerppola, 2008), recombinant construct pBiFCt-2in1-NC containing TTL3 and different interaction protein candidates TUB6, ACT2, CLC1 (for primers see Table S1) were constructed and transferred into *Agrobacterium tumefaciens* strain GV3101. These were transiently transformed into tobacco or onion leaf epidermal cells and placed back in the cultivation room. Fluorescence was observed on a confocal microscope (LSM880, Zeiss) 2 days after transformation. The vector pBiFCt-2in1-NC includes mRFP1 as a reference marker for transformation and expression.

## Results

The *TETRATRICOPETIDE-REPEAT THIOREDOXIN-LIKE 3* (*TTL3*) gene was identified from an independent collection of ~2,000 Mexican enhancer and gene trap lines (METs and MGTs) harboring Ds element with β-glucuronidase (GUS) reporter (Sundaresan et al., 1995; Estrada-Luna et al. 2002) which was screened for their promoter activity in the pericycle and developing LRPs. The gene trap line MGT182 showed strong activity in developing LRPs and lateral roots This pattern was identical to an independent construct described later. Using TAIL PCR (Liu and Whittier, 1995), T-DNA flanking sequence was identified as *TTL3* gene (AT2G42580).

### The *ttl3* affects the growth and development of lateral roots

To follow the role of *TTL3* in Arabidopsis root system development, seedling of *ttl3* and respective wildtypes were scanned, and morphometric traits of their respective root systems were recorded (Fig. 1). The total length of the *ttl3* root system is shorter at 8 and 9 DAG. Interestingly, this effect was not significant earlier (Fig. 1D), mostly due to the lack of developed lateral roots. The length of the primary roots was not significantly affected (Fig. 1E). The principal source of the difference was decreased total length and number of lateral roots in *ttl3* compared to wild type plants (Fig. 1F and 1G). To further uncover the source of the *ttl3* related decrease in the LRs, the total number of LR initiation events (including LRP) (Fig. 2A), the maximal LR length reached along acropetal sequence during the cultivation period (Fig. 2B), the length of root branching zone (Fig. 2C), and the LR branching density (Fig. 2D) were estimated. Among the measured parameters, *ttl3* had a significantly decreased density of emerged lateral roots (Fig. 2D), and there was no significant difference in the initiation density (Fig. 2A - 2C). The decreased density of LRs in *ttl3* lines may thus have resulted from a decelerated rate of LRP development after initiation but prior their emergence from the parent root.

**Figure 1.**
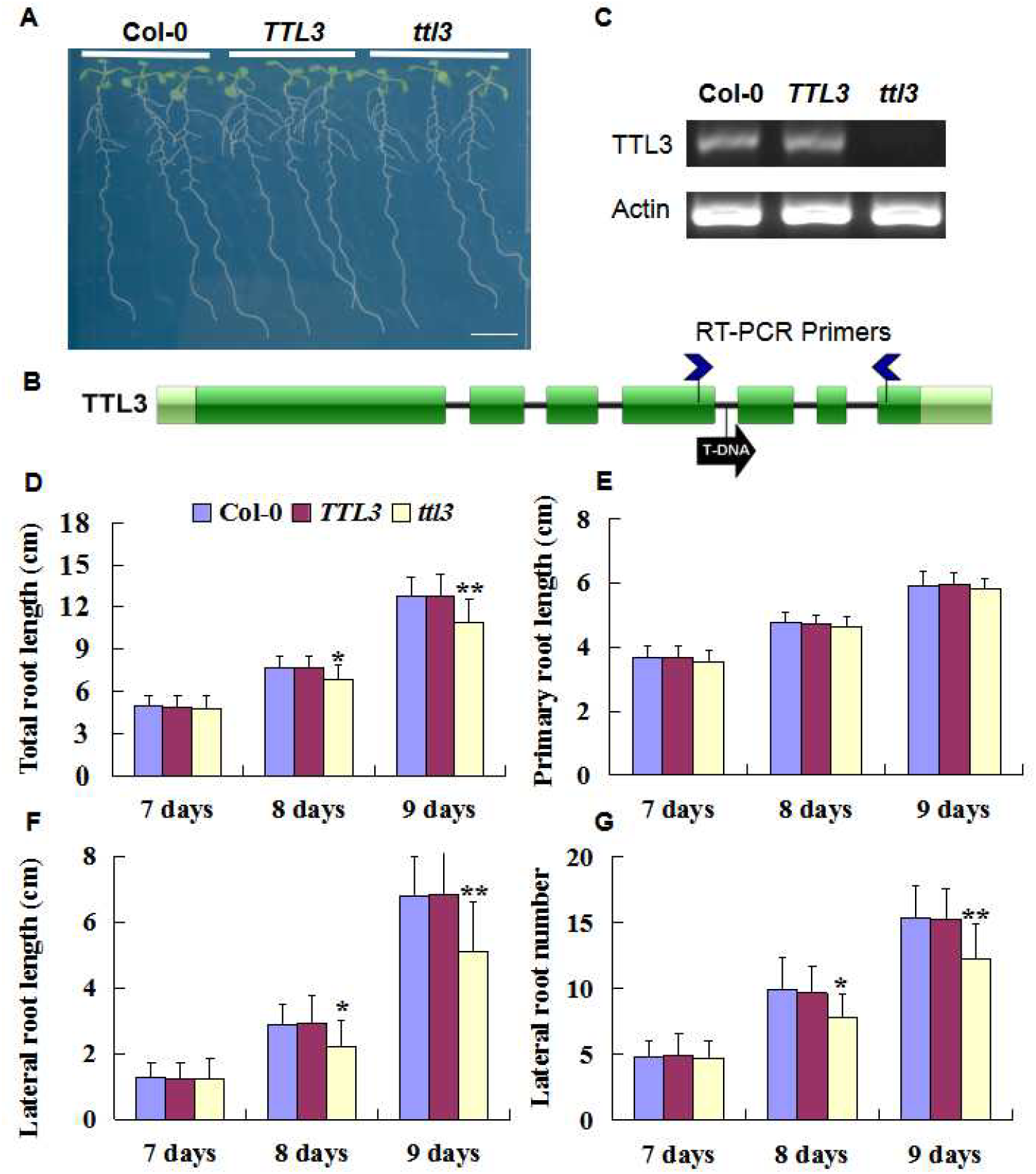
Phenotype of the *ttl3* mutant. (A) Seedlings of the Col-0, TTL3 and *ttl3*, 9 days after germination (DAG). Scale bar = 10 mm. (B-C) Transcript presence of TTL3 in Col-0, *TTL3* and *ttl3* by RT-PCR. and position of insertion in *ttl3* (D-G) Total root system length (D), primary root length (E), lateral root length (F) and lateral root number (G) of Col-0, *TTL3* and *ttl3* difference over time. Each data bar represents the means ± SE (*n* = 12). The asterisks indicate a significant difference from the corresponding control experiment by Student’s *t*-test (*: *p*< 0.05; **: *p* < 0.01).

**Figure 2.**
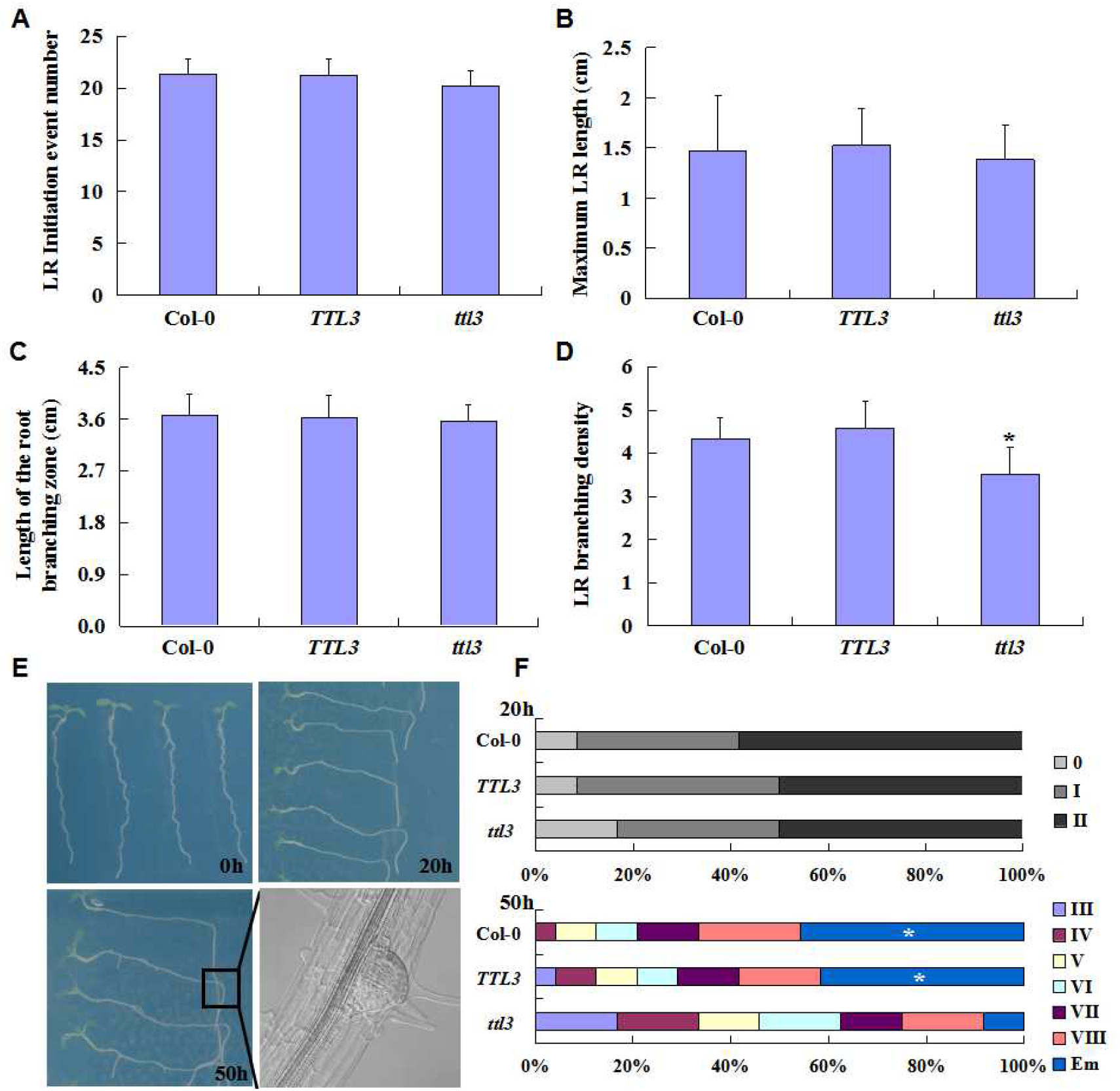
TTL3 affects lateral root development and emergence. (A-D) Total number of lateral root primordium (LRPs + LRs) initiation events (A), LR average length (B), Length of root branching zone (C), and LR branching density (D) were calculated 9 DAG. Each data bar represents the means ± SE (*n*=12). Asterisk indicates statistically significant differences between wide type and *ttl3* as determined by Student’s *t*-test (\**p* < 0.05; *n*=12). (E) Gravity stimulation of synchronised LRP. The 5-day-old seedlings were turned by 90° and grew for 20 and 50 h and then were analysed for stage of development of LRPs (F). The proportion of lateral root primordia at different stages of Col-0, *TTL3*, and *ttl3* after 20 hours and 50 hours (*n* = 24). The asterisks indicate a significant difference from the sequential pairwise comparisons by Pearson’s independent test (α = 0.05).

The pace of LRP development was further evaluated using a root bending test (Fig. 2E). LRPs were in a majority of roots induced at the bending point, 20 h after gravity-stimulus treatment (Fig. 2F).The induction rate was similar across all tested lines - Col-0, TTL3, and *ttl3*, confirming that the initiation and early development of LRP (from stage I to stage II) was not affected, which was in accord with *TTL3* expression described below. However, over 40% of the Col-0 and *TTL3* seedlings showed emerged lateral roots 50 h after gravitropic induction; this was a significantly higher proportion than the number of emerged lateral roots observed in *ttl3*, where only 10% of LRPs reached this point, indicating that the *ttl3* delays the emergence of LRP development (Fig. 2F). This effect was complemented once *pTTL3:TTL3-GFP* was transformed into *ttl3* background (*cTTL3*). The complemented line, *cTTL3*, exhibited wild-type traits regarding the number of LRs, as well as the timing of LRP development (Fig. S1). No obvious abnormalities of developing LRPs in either genetic background were observed.

**Figure S1.**
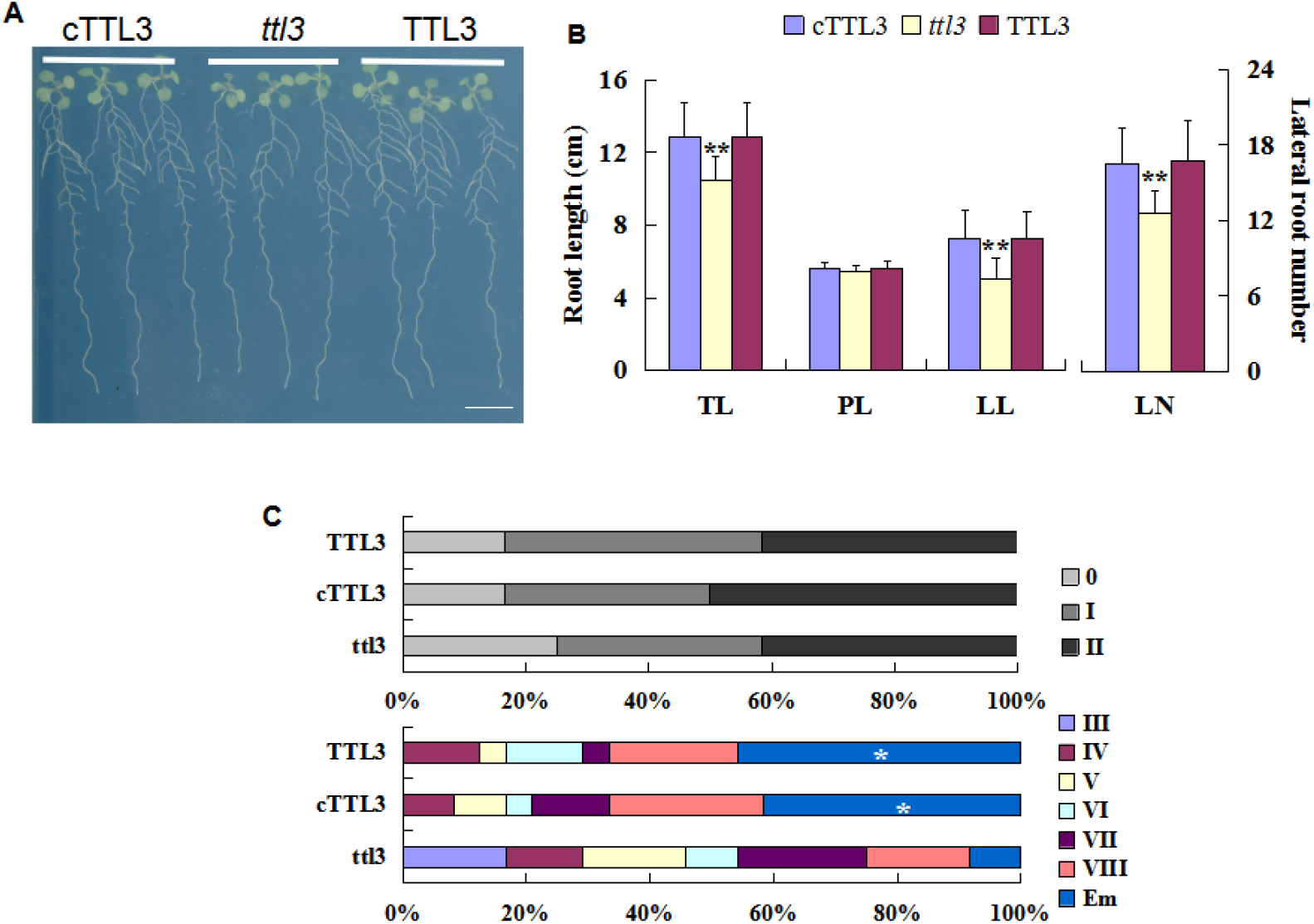
Phenotype of *ttl3* might be complemented. (A) Seedling of the cTTL3, *ttl3*, and TTL3 at 9 days after germination (DAG). Bar=10 mm. (B) Total root system length (TL), primary root length (PL), lateral root length (LL) and lateral root number (LN) of cTTL3, *ttl3*, and TTL3 9 days after germination (DAG). Each data bar represents the means ± SE (*n* = 12). The asterisks indicate a significant difference from the corresponding control experiment by Student’s *t*-test (*: *p*< 0.05; **: *p* < 0.01). (C) Root bending assay. The proportions of *cTTL3, ttl3, and TTL3* lateral root primordia at different stages were measured 20 and 50 hours (*n* = 12; 24) later. The asterisks indicate a significant difference from the sequential pairwise comparisons by Pearson’s independency test (α = 0.05).

To test for additive/redundant effects of other TTL genes, the triple mutant *ttl1ttl3ttl4* was constructed and phenotype described above was re-evaluated (Fig. 3A - 3C). The root system, including primary root and lateral root, was more affected in the triple mutant (Fig. 3D). This stronger phenotype was used to search for processes modifying the growth of roots. The observed change in growth might be either related to cell elongation, their proliferation or both. The origin of the *ttl1ttl3ttl4* phenotype can be seen in the length of the meristematic zone (Fig. 3E and Fig. 3F), rather than the length of differentiated cells (Fig. 3G). The cell proliferation assay (Fig. 3C) demonstrates that the phenotype is associated primarily with RAM activity (Fig. 3H).

**Figure 3.**
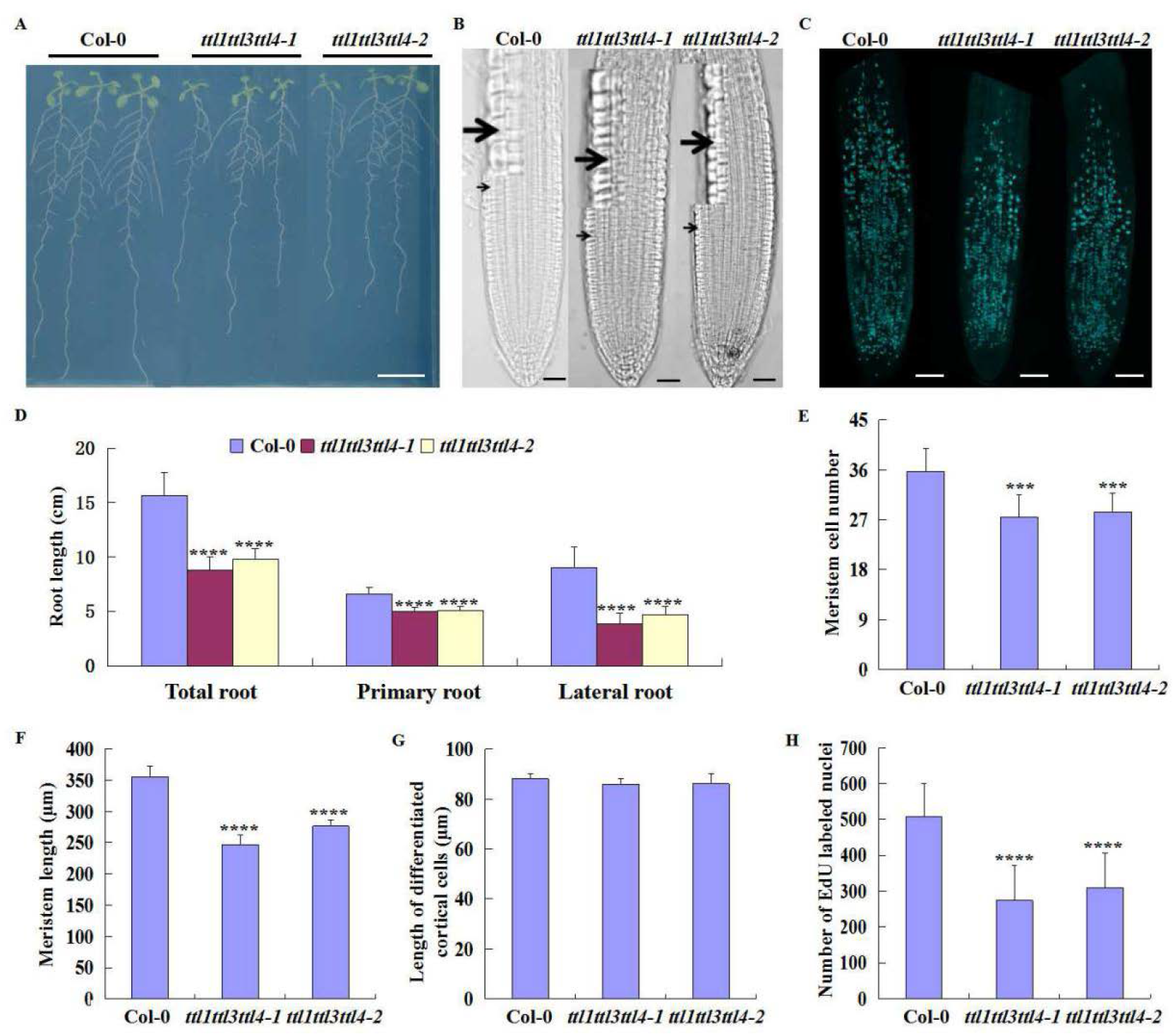
Phenotype of *ttl1ttl3ttl4* mutant. (A) Seedling of the Col-0, *ttl1ttl3ttl4-1*, and *ttl1ttl3ttl4-2* 9 days after germination (DAG). Scale bar = 10 mm. (B) Length of the meristematic zone in the Col-0, *ttl1ttl3ttl4-1*, and *ttl1ttl3ttl4-2* at 7 DAG. The enlarged inlay displays detail of the edge between elongation and transition zone. Scale bars = 50 μm. (C) The distribution of dividing cells in the primary root meristem of Col-0, *ttl1ttl3ttl4-1*, and *ttl1ttl3ttl4-2* at 7 DAG shown on the 20 μm z-stacks MAX projection. Scale bars = 50 μm. (D) Data analysis of total root length, primary root length, and lateral root length of Col-0, *ttl1ttl3ttl4-1* and *ttl1ttl3ttl4-2*. (E-F) The number of cells in the meristem (E) and meristem length (F) of the primary root in Col-0, *ttl1ttl3ttl4-1*, and *ttl1ttl3ttl4-2*. (G) Analysis of the length of differentiated cortical cells in Col-0, *ttl1ttl3ttl4-1* and *ttl1ttl3ttl4-2*. (H) The number of S-phase cell nuclei in the primary root apical meristem labelled with EdU in the Col-0, *ttl1ttl3ttl4-1*, and *ttl1ttl3ttl4-2*. Each data bar represents the means ± SE, *n* = 10 (E, F, G), n≥22 (H). The asterisks indicate a significant difference between genotypes evaluated by Student’s *t*-test (***: *p* < 0.001; ****: *p* < 0.0001).

It was reported that the TTL protein scaffolds brassinosteroid signalling components and root growth of the *ttl3* mutants are hyposensitive to BR (Amorim-Silva et al., 2019). Therefore, we analyzed the effect of BR on the development of LRPs for *ttl3* and its respective WT (Fig. 4A). After 6 days of 100 nM eBL treatment, the length of the root branching zone for *ttl3* was not significantly different from *TTL3*. However, the change in the number and density of LRs in *ttl3* mutants were less affected compared to *TTL3* plants. In the *TTL3* plants, the number of LRs reached 61% of non-BR treatment, and in the *ttl3* plants, the proportion was 75%, which was significantly higher than that of the *TTL3* plants (Fig. 4C). Also, the density of LRs of the *ttl3* mutants accounted for 97% of non-BR treatment, which is higher than TTL3 plants (88%) (Fig. 4D). LRP development of the eBL-treated roots was delayed in *TTL3* plants. However, there was no further significant decrease in percentage of LRs in *ttl3* plants (Fig. 4E).

**Figure 4.**
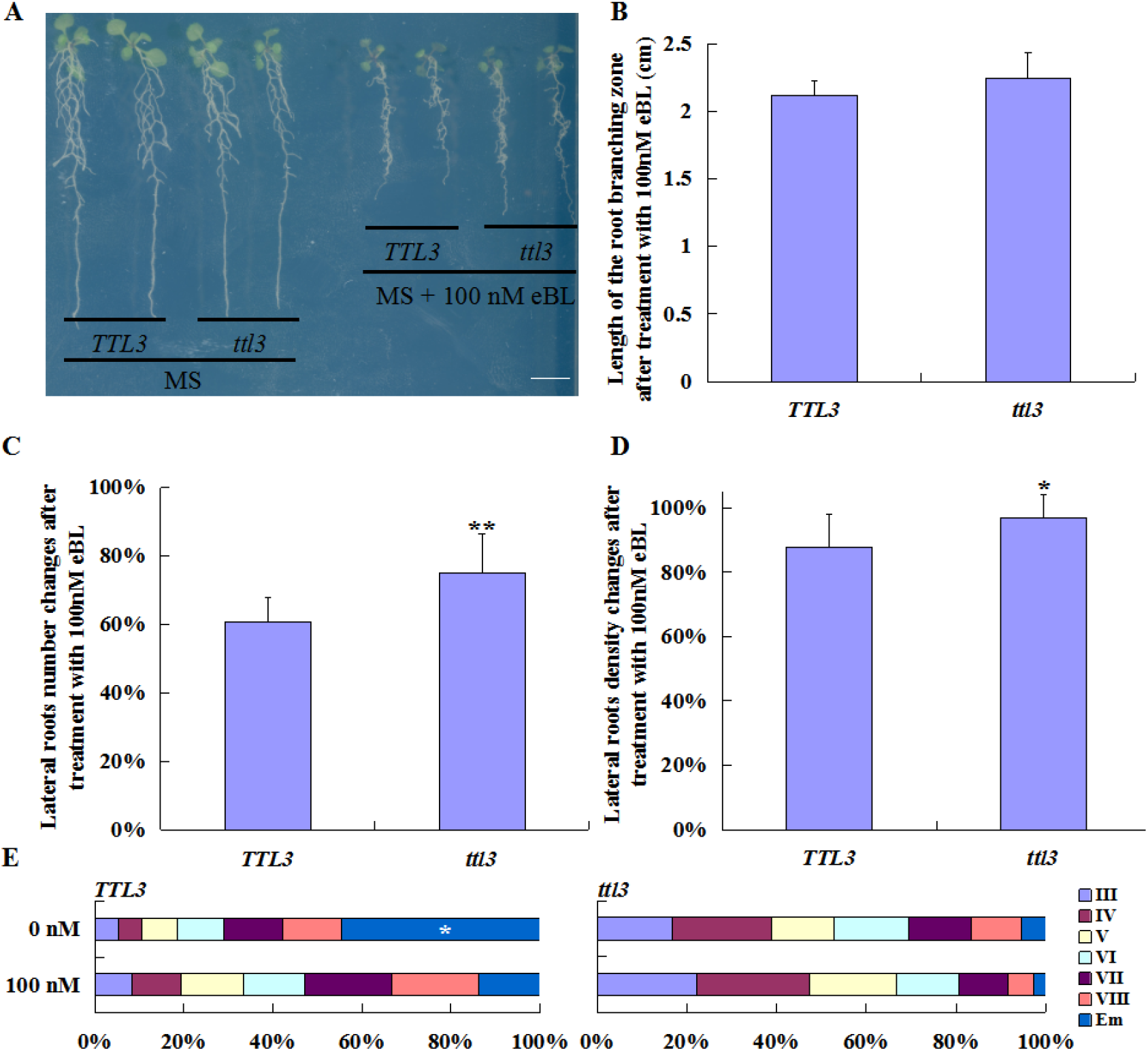
The *ttl3* mutants are less sensitive to BR treatment. (A) Seedlings were grown for 5 days in 0.2× MS medium and then transferred to 0.2× MS medium supplemented with 100 nM eBL, photographs show representative seedlings after 6 days 100 nM eBL treatment. Scale bar = 10 mm. Length of lateral root branching zone (B), percentage of lateral root number (C) and the lateral root density (D) in *TTL3* and *ttl3* after 6 days of 100 nM eBL treatment. Asterisks indicate statistically significant differences between *TTL3* and *ttl3* as determined by Student’s *t*-test (*: *p*< 0.05; \*\**p* < 0.01; *n*=12). (E) Root bending synchronised LRPs (after 50 h) indicate the effect of 100 nM eBL on the development of *TTL3* and *ttl3* lateral root primordia (*n*≥36). The asterisks show a significant difference from the sequential pairwise comparisons by Pearson’s independent test (α = 0.05).

### Expression pattern of *TTL3* in Arabidopsis lateral root

Promoter activity of *TTL3* was detected in both primary and lateral roots. *TTL3* transcription takes place in the protodermal tier of the meristematic zone of the primary root (Fig. 5E). A stronger response was found in the RAM of growing LRs (Fig. 5D) and their primordia from the stage I (Fig. 5A and 5B), through emergence and later in the RAM of emerged LR (Fig. 5C and 5D). However, the *TTL3* promoter activity was not present before and during LRP initiation. *TTL3* expression was also found in developing leaf venation (Fig. 5F). The expression pattern of *TTL3* was almost complementary to *TTL1* and *TTL4* in roots. *TTL1* (Fig. S2) starts to be transcribed within the procambial strands at transition domain of the RAM (Ivanov and Dubrovsky, 2013). It is obvious from more differentiated positions that it is not xylem, but a complementary, phloem associated part of the central cylinder. Weaker promotor activity was also found in the protoderm of the transition domain of the RAM and early elongation zone. In leaves, *TTL1* expression is associated with vasculature tissues and stomatal guard cell differentiation. *TTL 4* is transcribed in the central cylinder of both primary and lateral roots. In growing roots, it normally starts in the transition domain of the RAM and elongation zone and disappears in fully differentiated tissues. *TTL4* transcription takes place also during differentiation of leaf mesophyll and disappears in mature tissues (Fig. S2). None of the TTL1, 3, or 4 were present in the pericycle, founder cells or stage I LRPs.

**Figure 5.**
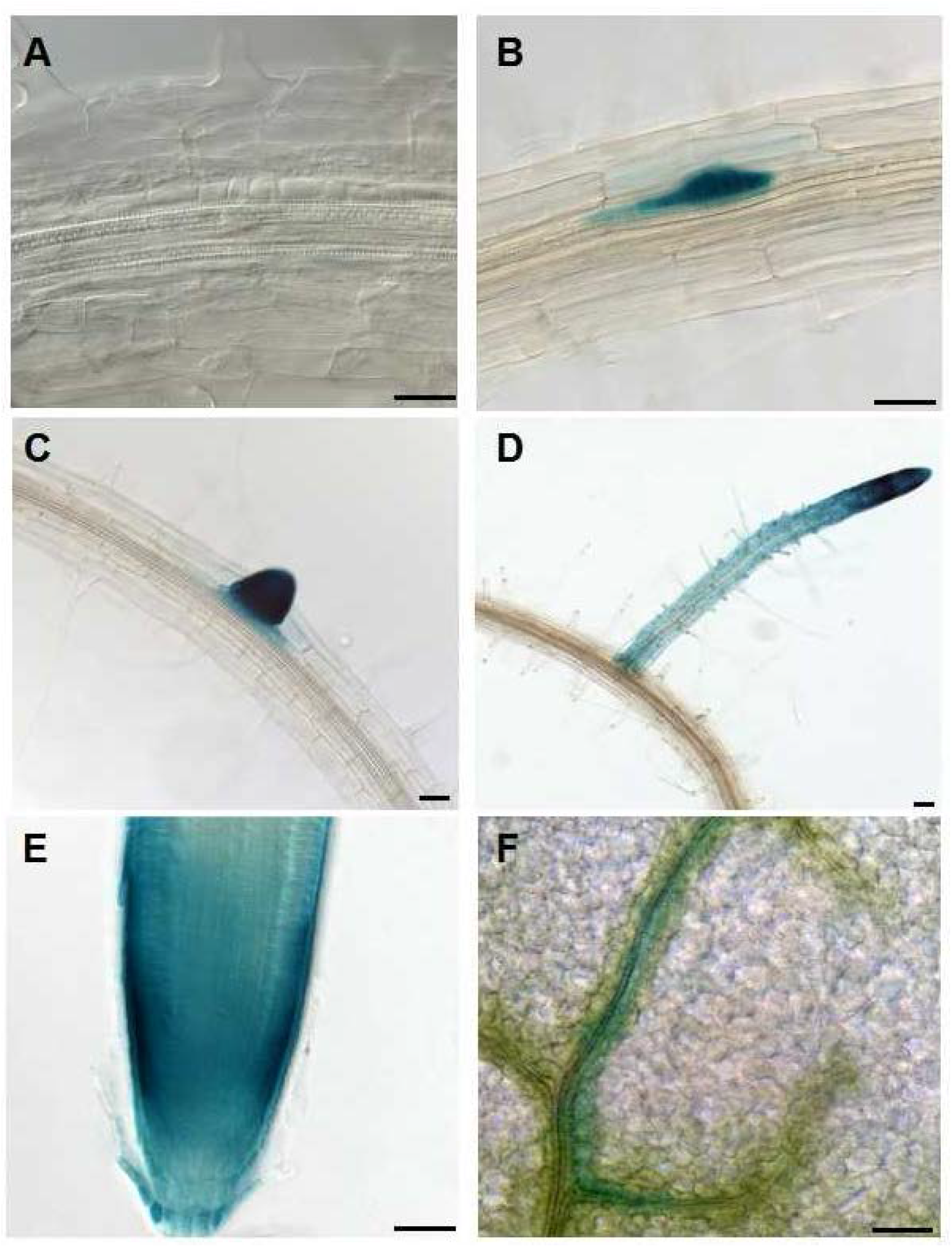
The *TTL3* promoter activity monitored by *pTTL3-GUS* transgene. Promotor activity is not present in the earliest stages of lateral root primordium (A), and appears in later stages of lateral root primordium (B), recently emerged lateral root (C), and growing lateral root with promotor activity in cortex and outer tiers of root apex (D), primary root (E), and developing leaf vein (F). Scale bars = 50 μm.

**Figure S2.**
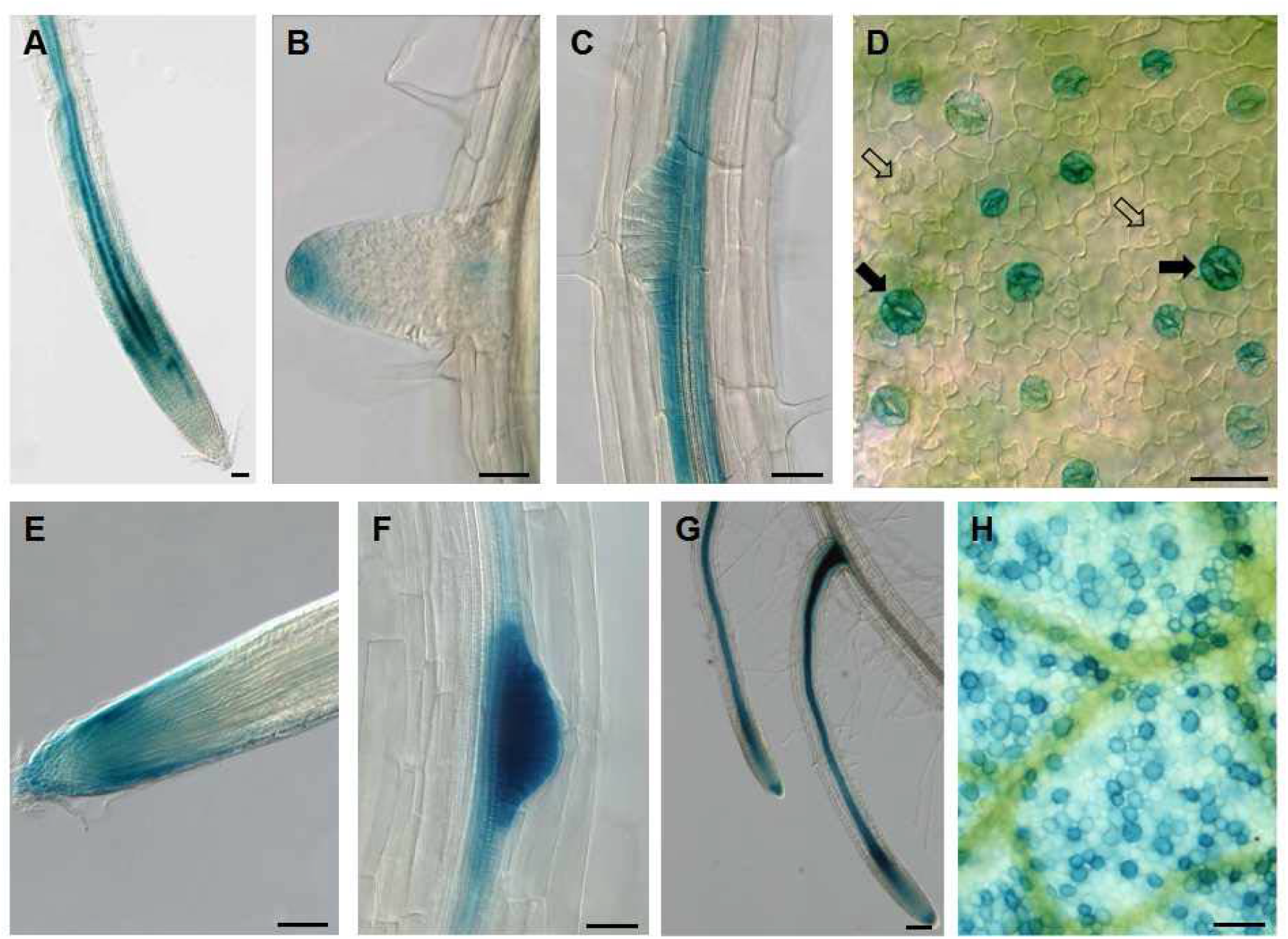
The promoter activity of *TTL1* and *TTL4*. *pTTL1-GUS* expression in the central cylinder of primary root (A), early lateral root (B), lateral root primordium (C) and differentiating guard cells (D; full arrowheads = guard cells, empty arrowheads = meristemoids). *pTTL4-GUS* expression in the primary root tip (E), lateral root primordium (E), positive central cylinder in growing lateral roots (F) and scattered cell in developing leaf mesophyll (H). Scale bars = 50 μm.

Transient transformation of tobacco (*Nicotiana benthamiana*) leaves with *35S::TTL3-GFP*. was used to follow cellular localization of the protein. Despite cytoplasmic localization of *TTL3*, the dominant signal was detected in a network structure of cytoskeleton (Fig. 6B), which is lost if microtubules are chemically depolymerized (Fig. 6C - 1mM oryzalin). High affinity of TTL3-GFP was indicated by rapid labelling of growing microtubules (Video S1). Moreover, some bead-like structures were found to move along the microtubular network (Fig. 6A; Video S1).

**Figure 6.**
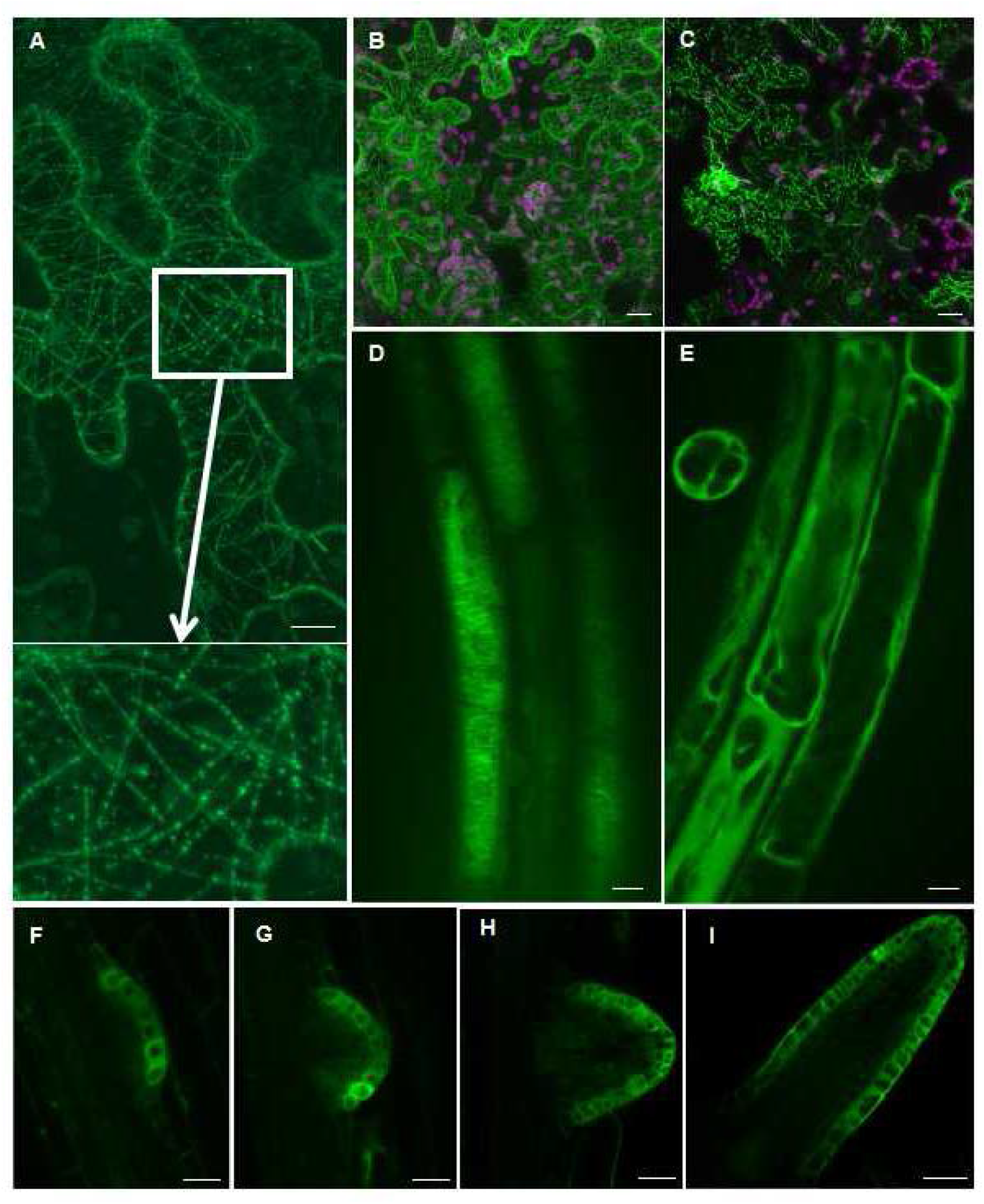
Subcellular localization of TTL3. (A-B) High affinity of *35S::TTL3-GFP* to microtubules were observed in tobacco leaves (*N. benthamiana*) cells. The enlarged inlay shows a portion of the white box with clear microtubule-related signals, and there are beaded signals on the microtubules. Scale bars = 30 μm. (C) Oryzalin (1mM) treatment disrupted the microtubules and pattern of TTL3-GFP distribution. Scale bar = 30 μm. (D) Cortical microtubule signals can be observed in the root elongation zone of the *pTTL3::TTL3-GFP* Arabidopsis. Scale bar = 10 μm. (E) The *pTTL3::TTL3-GFP* Arabidopsis line was treated with Oryzalin (100 μM), and observed 10 minutes later. Scale bar = 10 μm. (F-I) *TTL3* is expressed in the outermost layer of lateral root primordium at different stages. Scale bars = 20 μm.

The Arabidopsis (Col-0 and *rdr6* background) plants containing *pTTL3::TTL3-GFP* allowed for the localization of *TTL3* expression *in planta*. The network pattern typical of cortical microtubule signal and cytoplasmic signal were found in the elongation zone of the primary root (Fig. 6D; Video S2). This microtubule associated pattern disappeared after the oryzalin (100 μM) treatment (Fig. 6E; Video S3). Consistent with the *pTTL3-GUS* pattern, the protein was never presented during LRP initiation but appeared during early stages of LRP development (approximately from stage II) and remained active during later development (Fig. 6F and 6G; Fig. 7A), LRP emergence and growth of lateral roots (Fig. 6H and 6I; Fig. 7B-7D). Within the developing LRPs, the TTL3 was expressed in the outermost layer of the central domain of LRPs. This localization appeared usually during the stage II and was maintained during the emergence and development of lateral roots. Similarly, the bead-like structure was found in the differentiated cells of the Arabidopsis root (Video S4) where expression and cytoplasmic background was significantly lowered.

**Figure 7.**
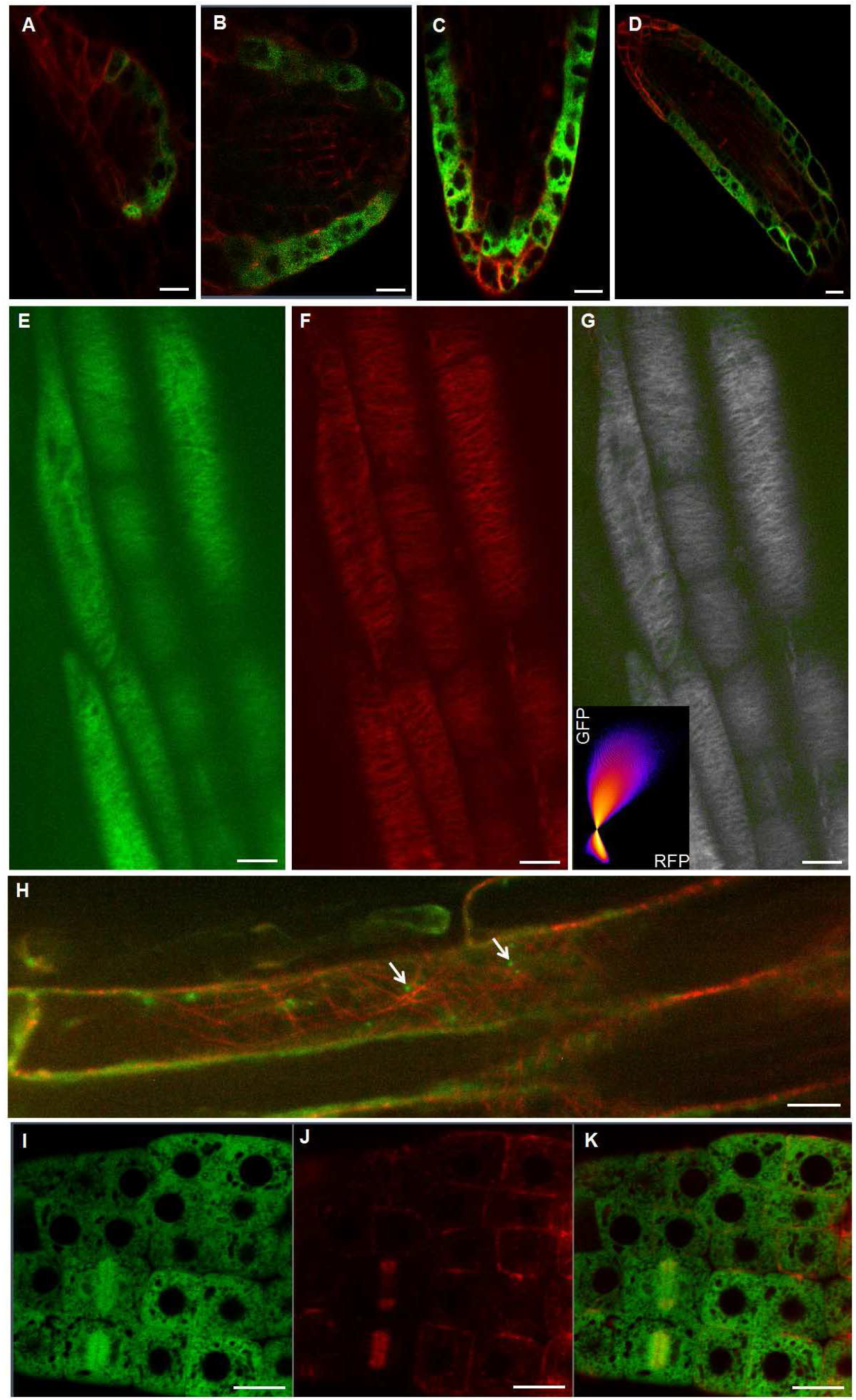
TTL3 is associated with microtubules in the roots of Arabidopsis. (A-D) The different stages of lateral root development in plants harbouring both *pTTL3::TTL3-GFP* and *35S::MAP4-RFP*. Scale bars = 20 μm. (E-G) TTL3 and cortical microtubules colocalize in the primary root elongation zone. *pTTL3::TTL3-GFP* (E) *35S::MAP4-RFP* (F), and merged (G) images are shown. The colocalization graph is displayed in the lower-left corner of the merged image (G): Manders’ tM1 (above the automatic threshold of GFP) is 0.860, and Manders’ tM2 (above the automatic threshold of RFP) is 0.816. Scale bars = 10 μm. (H) TTL3 moved along microtubules in *pTTL3::TTL3-GFP* transgenic Arabidopsis differentiated cells. The arrows pointed to the TTL3 moving signal. Scale bar = 10 μm. (I-K) TTL3 colocalized with cortical microtubules (MTs) in the phragmoplast of lateral root meristematic cells. *pTTL3::TTL3-GFP* (K), *35S::MAP4-RFP* (L), and merged (M) images are shown. Scale bars = 10 μm.

Cellular localization of TTL3-GFP in Arabidopsis roots varied according to position along the root. We focused on lateral roots, where the signal was stronger and easier to follow. In meristematic cells, the localization of TTL3-GFP was rather intense and cytosolic background did not allow for distinction of specific localization within the cells. This situation was valid for both LRPs and LRs. However, within the transition domain of the RAM of LRs, the TTL-GFP protein decorated cortical microtubules and this pattern remained within the protoderm till the late elongation zone (Fig. 6D). The spatial association of TTL3 protein and microtubules was rather obvious from transformants containing both *35S::MAP4-RFP* and *pTTL3::TTL3-GFP* (Fig. 7E-7G; Video S5). In protodermal cells of the elongation zone, the TTL3-GFP signal appeared in bead-like bodies moving along cortical microtubules (Fig. 7H; Video S6). Interestingly, TTL3 was localised also in the phragmoplast of dividing cells within the RAM (Fig. 7I-7K). TTL3 presence detectable in the meristematic, transition and elongation zone disappears with the advancement of differentiation. As might be expected from the Western blot (Fig. S4) part of the cytoplasmic signal should be attributed to presence of free GFP, formed by partial cleavage of fusion protein.

### Searching for TTL3 interacting proteins

To further understand the function of TTL3 protein in the lateral root development, we performed co-immunoprecipitation screening for interactive partners in Arabidopsis. Total protein extracts from *pTTL3::TTL3-GFP* and *35S-GFP* root systems were incubated with GFP-Trap-A agarose beads to pull down TTL3-GFP. Free GFP was used as a nonspecific control to subtract non-specific candidates from the output. There were 421 candidate proteins that putatively interacted with TTL3, 259 of which were categorized into GO PANTHER Protein Classes. The annotated proteins covered 17 protein classes (Fig. S3).

These 17 classes can be roughly divided into 5 categories (Table S2): Enzyme (133, 51.4%), Nucleic Acid Binding (56, 21.6%), Transport protein (35, 13.5%), Cytoskeleton protein (23, 8.9%), others (12, 4.6%). Of these categories, transport and cytoskeleton protein are most consistent with the intracellular localization of the TTL3 protein. Among the enzymes, there are 25 transferases, plus 35 transport proteins (17 transporters, 16 membrane traffic proteins, 2 carrier proteins), and 60 interaction proteins are related to intracellular transport. The cytoskeleton proteins include 11 tubulins, 6 actins, 6 cytoskeleton related proteins (Table S2).

**Table S2.** Categories of TTL3 interaction partners separated by co-immunoprecipitation

| Categories | Quantity | Detailed classification (corresponding number) |
| --- | --- | --- |
| Enzyme | 133; 51.4% | Hydrolase (38), Oxidoreductase (33), Transferase (25), Enzyme Modulator (19), Ligase (9), Isomerase (5), Lyase (4) |
| Nucleic Acid Binding | 56; 21.6% | Nucleic Acid Binding (56) |
| Transport protein | 35; 13.5% | transporters (17), membrane traffic proteins (16), carrier proteins (2) |
| Cytoskeleton protein | 23; 8.9% | tubulins (11), actins (6), cytoskeleton related proteins (6) |
| Others | 12; 4.6% | Chaperone (6), Transcription factor (3), Adaptor Protein (1), Cell Junction Protein (1), Immunity Protein (1) |
| Total | 259 |  |

**Figure S3.**
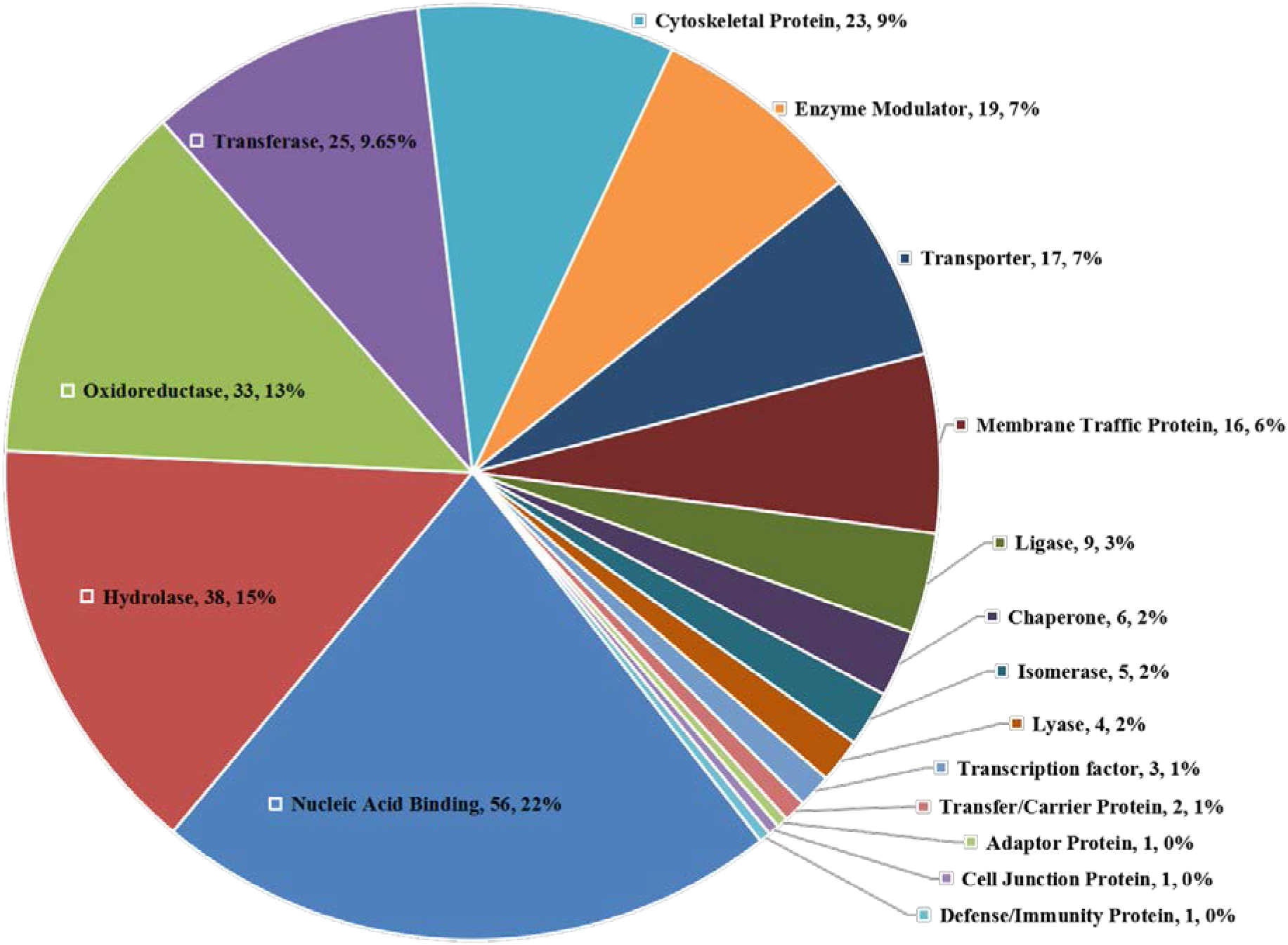
Detailed classification of TTL3 interacting partners separated by co-immunoprecipitation. The classification according to PANTHER Protein Class.

According to the Co-IP results and the characteristics of intracellular localization of TTL3 protein, we chose CO-IP candidates ACT2 (AT3G18780), TUB6 (AT5G12250), and CLC1 (AT2G20760) to confirm their physical interaction via a yeast two hybrid assay (Y2H). TTL3 was fused to the pGADT7 vector and the candidate proteins (ACT2, TUB6, and CLC1) were fused to the pGBKT7 vector. As shown in Fig. 8A, TTL3 direct interaction was confirmed only with TUB6 in the Y2H. Bimolecular Fluorescence Complementation (BiFC) was employed to further support the expected interaction. A recombinant construct pBiFCt-2in1-NC containing TTL3 and different candidate interacting proteins (TUB6, ACT2, CLC1) were transfected into the tobacco or onion epidermal cells. YFP fluorescence signal indicative of physical interaction between putative partners was found only in the combination of TUB6 and TTL3 proteins (Fig. 8B-8E) but not the others. These results indicate that TTL3 directly interacts with TUB6 *in vivo*.

**Figure 8.**
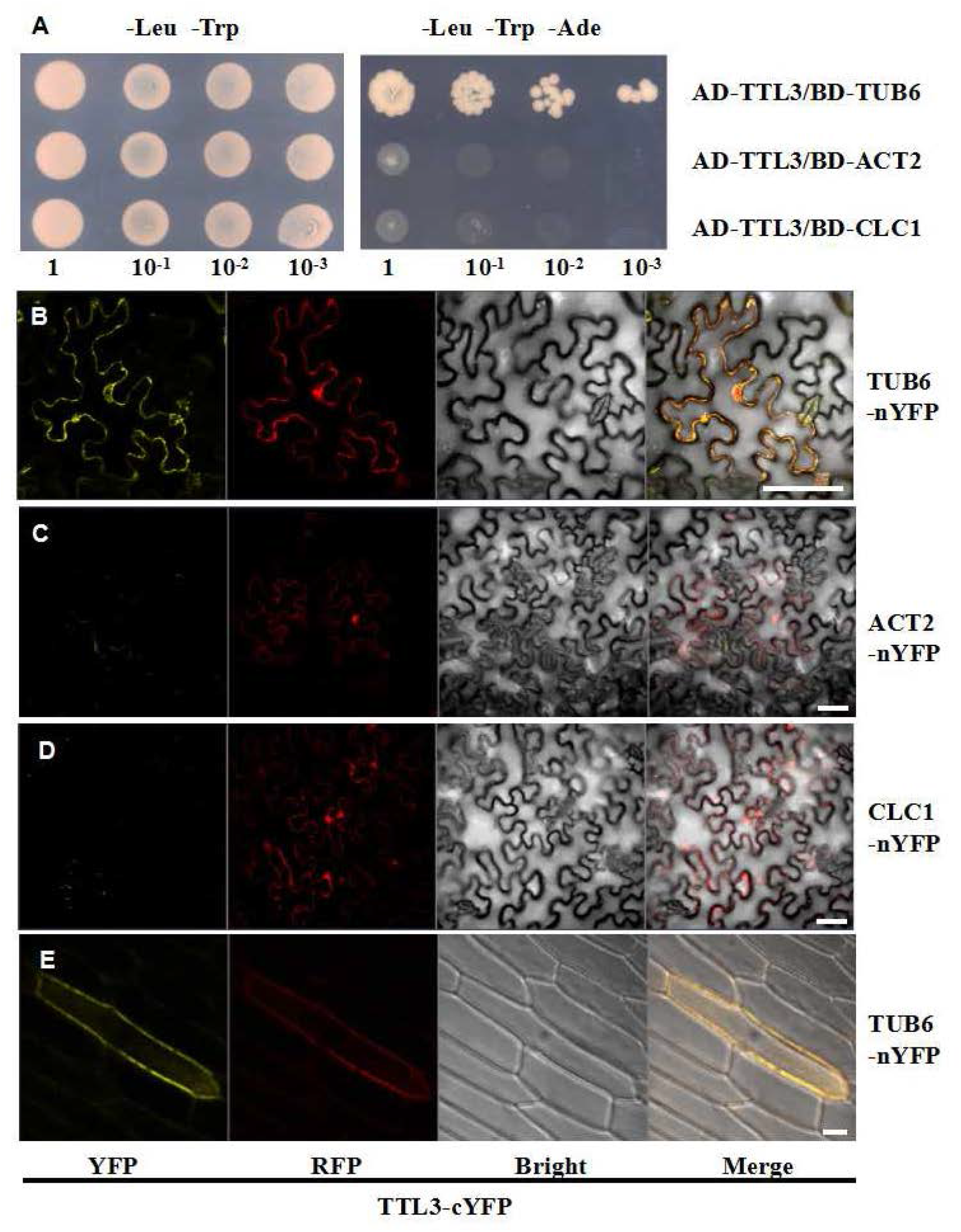
TTL3 interacts with TUB6. (A) Yeast two hybrid assay to determine the interaction of TTL3 with TUB6, ACT2 and CLC1. Growth on plasmid-selective media (left column) and interaction-selective media (lacking adenine; right column) are shown. (B-E) BiFC-based analysis of the TTL3 and candidate interaction proteins in transiently transformed tobacco (B-D) and onion (E) epidermal cells. Representative images of laser scanning confocal microscopy of tobacco leaf transfected with *Agrobacterium tumefaciens* either containing TTL3/TUB6 (B), TTL3/ACT2 (C), TTL3/CLC1 (D), and of onion epidermal cells transfected with *Agrobacterium tumefaciens* containing TTL3/TUB6 (E). YFP fluorescence, RFP fluorescence, bright filed and merged images are shown for each transformation combination. Scale bars = 50 μm.

**Figure S4.**
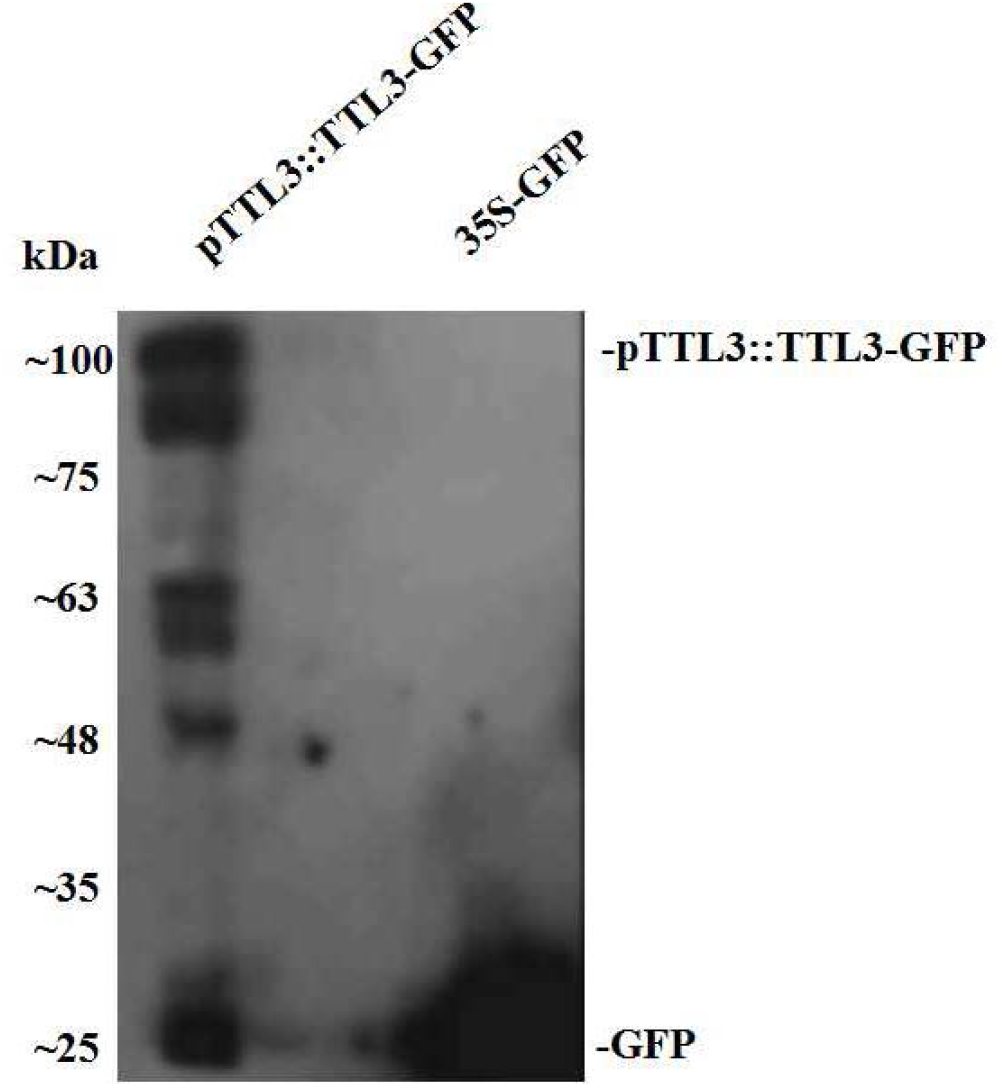
Western blot analysis for the *pTTL3::TTL3-GFP* and *35S-GFP* in Arabidopsis transgenic plants. Total proteins were extracted from 2-week-old seedlings of *pTTL3::TTL3-GFP* and *35S-GFP* lines, followed by western blot analysis using GFP antibodies.

## Discussion

Roots evolved as a specialized organ for interacting with a heterogeneous edaphic environment (De-Jesús-García et al., 2020). Its architecture not only allows for efficient exploration of terrestrial resources, but also affects the economy of energy and biomass investment, which is crucial for plant fitness and survival under stress conditions. Lateral roots contribute the majority of root biomass and rhizosphere interface (Dittmer, 1937). Our knowledge of LR development, and their regulatory and executive networks has increased tremendously during the last two decades. However, its complexity still provides vast space for research and improvements of our understanding.

In our search for novel regulatory elements influencing lateral root development, we identified TTL3 (Tetratricopeptide-repeat Thioredoxin-Like 3, AT2G42580), which was found to be involved in osmotic stress response (Lakhssassi et al., 2012). TTL3 is one of four members of a gene family in Arabidopsis. Characteristic combination of three tandem tetratricopeptide domains and thioredoxin-like domain is specific only for terrestrial plants (Rosado et al., 2006; Lakhssassi et al., 2012). The functional role of the TTL3 protein in lateral root development is still rather unclear despite evidence that it is related to the stress response (Rosado et al., 2006; Lakhssassi et al., 2012) and differentiation of leaf vascular tissues (Ceserani et al., 2009). Physical interaction of TTL3 with VH1/BRL2 receptor-like kinase (Ceserani et al., 2009), BRASSINOSTEROID INSENSITIVE1 (BRI1) receptor kinase, BRI1-SUPPRESSOR1 (BSU1) phosphatase and the BRASSINAZOLE RESISTANT1 (BZR1) transcription factor, suggest its involvement in brassinosteroid signaling (Amorim-Silva et al., 2019). Modification of primary root growth-sensitivity to BRs in single and multiple TTL knock-out mutants is in accordance with this conclusion and also indicates possible functional redundancy within the gene family (Amorim-Silva et al., 2019). In our results, the number and density of LRs in *ttl3* mutants were less affected compared to wild-type plants after 6 days of 100 nM eBL treatment (Fig. 4), indicating that this trait of root architecture is under BR control and that TTL3 is a part of the circuit. Regarding the functions of other TTL family members, TTL1 is most likely involved in ABA sensitivity as the *ttl1* mutant is more sensitive to osmotic stress (Rosado et al., 2006). TTL2 takes part in male gametophyte development. In plants with multiple TTL gene mutations, primary root growth was decreased under osmotic stress and BR treatment (Lakhssassi et al., 2012). The promoter activity localization of TTL1 and TTL4 are almost complementary to the TTL3, which is active in developing procambium, where TTL3 is not found, of the primary and lateral roots (Fig. 5 and S2). Also interesting is that their expression in above-ground tissues, where TTL1 and TTL4 are involved in differentiation of leaf mesophyll, is particularly conspicuous in the substomatal cavity in the hypocotyl (TTL4 - Fig. S2) or during differentiation of stomatal guard cells and the formation of the stomatal aperture (TTL1 - Fig. S2). The expression pattern is bound to meristems and differentiating tissues, which localizes their role to associated processes. However, such a conclusion might be questioned by a report that TTL3 mRNA might be mobile in vascular tissues (Thieme et al,.2015).

Primary root growth is not distinctly affected in *ttl3* mutant plants (Amorim-Silva et al., 2019 and this study), however the principal effect in root system was pronounced in the development of lateral roots. Searching for the source of the *ttl3* phenotype we concluded that it is not the initiation of LRPs, as is in accordance with the expression pattern of TTL3, but later during the development of the primordia and after root emergence. The TTL3 protein was detected in LRPs from stage II and later (Fig. 5G-5J). Particularly, the timing of pre-emergent development was affected as is estimated from synchronized LRP initiation after primary root bending. In this experiment, a significantly higher proportion of LRPs develop to later stages of LRP and emerged lateral roots. Therefore, we have found a stable phenotype of the *ttl3* knock out mutant affecting root system development during the post-initiation stage of lateral root development with a pronounced effect on the timing of LR development.

### TTL3 affects lateral root emergence and development

LR development is a complex, stepwise process (Banda et al. 2019; Torres-Martinez et al. 2019), regulated by a network of diverse factors. Complex biomechanical and biochemical interactions between LRP and the surrounding tissues are involved in all the steps of LRP formation – from initiation, to growth, to emergence (Vilches-Barro and Maizel, 2015; Stoeckle et al., 2018). Auxin is believed to play a coordinating role in multiple stages throughout LR development. It attunes initiation and spatial distribution of LRP; regulates nuclear migration in LRP founder cells; influences interactions between LRP and their overlying tissues, and facilitates penetration of emerging LRP (Du and Scheres, 2018). LRP growth is commonly divided into seven stages: stage I to stage IV of the LRP occur bound by the endodermis, while stages V through VII take place after endodermal penetration (Malamy and Benfey, 1997). Before the emergence and formation of a lateral root, the LRP must cross three cell layers: the endodermis, the cortex, and the epidermis (Stoeckle et al., 2018). When LRP form four cell layers (stage IV), stem cells are defined within arising apical meristem, and the LRP breaks through the endodermis at approximately the same time (Banda et al. 2019). Auxin acts as a local signal released by LRP to facilitate reprogramming of overlying tissues (Swarup et al., 2008), including modification of their cell walls and their hydraulic properties (Péret et al., 2012; Roycewicz and Malamy, 2014). Auxin (IAA) can enter cortical cells to induce LAX3 expression, which creates a positive feedback loop that triggers high auxin levels and subsequently induces cell wall remodeling (CWR) genes, such as polygalacturonases (Swarup et al., 2008). Auxin inhibits the expression of almost every member of the aquaporins, which encode local water channel in the plasma membrane (PIPs) or vacuole membrane (TIPs) and regulate the hydraulic properties of plant cells (Péret et al., 2012).

This period of pre-emergent LRP development relates to TTL3 expression. During the development of LRP, TTL3 mainly is expressed in the outer-most layer (Fig 6G - 6I; Fig. 7A - 7B), which may be related to internal LRP development as well as to the communication between LRP and the overlying tissue. Experimental synchronization of LRPs further confirms that *ttl3* is affecting the LRP development before its emergence. The total length and number of emerged lateral roots in *ttl3*, which are significantly reduced compared to wild type plants at 8 and 9 days after germination (Fig. 1), thus results particularly from this pre-emergent and early post-emergent developmental phase. In addition, the maximum lateral root length (Fig. 2B) and the average lateral root length (data not shown) did not differ significantly between wild type and *ttl3* mutant. These results indicate the *ttl3* mutants hinder LRP development mostly before emergence from the overlying tissue but not during their later growth.

TTL3 participates in the BR signal transduction pathway (Amorim-Silva et al, 2019), which can trigger and modulate various developmental processes such as cell elongation, vascular differentiation, root protophloem development among others (Szekeres et al., 1996; Azpiroz et al., 1998; Catterou et al., 2001a; Catterou et al., 2001b; Kang et al., 2017). The BR-insensitive mutant *bri1* shows a dramatically decreased number of lateral roots. Low concentrations of BL promote the initiation of lateral roots, most likely via increased acropetal auxin transport (Bao et al., 2004). On the other hand, higher concentrations of BL suppress lateral root formation, possibly by inducing the expression of AUX/IAAs to inhibit auxin signaling (Gupta et al., 2015). BR can trigger rapid cell wall expansion in root cells before cell elongation, which is attributed to the loosening of the cell wall, and the absorption of apoplastic water. It depends on the activated proton pump of the plasma membrane (AUTOINHIBITED PLASMA MEMBRANE H^+^-ATPases), such as AHA1 which interacts with BRI1 (Caesar et al., 2011). A new study reveals that the main effect of BR signalling is on the distribution of auxin in the cell, which has a major impact on nuclear auxin signaling (Rana and Hardtke, 2020). The BR signal controls PIN-LIKE-dependent nuclear abundance of auxin (Sun et al., 2020). From this very brief overview it is clear that BR interaction with LRP development might be rather diverse and does not reasonably narrow the options of TTL3 functions. Searching for more sources of decreased root growth, which was more significant in the triple mutant *ttl1ttl3ttl4*, we tested the effect on elongation of cortical cells, which was not significant, and cell production in RAM. Primary RAM of the *ttl1ttl3ttl4* triple mutant were significantly shorter, containing fewer meristematic cells. And in the EdU-based proliferation assay, the number of S-phase nuclei labeling in the triple mutants was significantly lower than that of wild-type plants (Fig. 3). In addition, TTL3 was enriched in the phragmoplast during cell division (Fig. 7I-7K). All the above indicates that the TTL3 might play a role in cell division and/or cells differentiation associated processes.

### TTL3 and its interacting partners

The TTL gene family is specific for terrestrial plants, with a unique combination of tetratricopeptide (TPR) and thioredoxin-like (TRXL) domains. TPR defines a loosely conserved 34 amino acids protein-protein interaction domain (Blatch and Lässle, 1999), which indicates a chaperone-like function of TTL proteins. In plants, TPR domains and their scaffold complexes are involved in various phytohormone responses, root development, plastid distribution and photosynthetic machinery (Jacobsen et al., 1996; Rosado et al., 2006; Yang et al., 2011; Hu et al., 2014; Bhuiyan Et al., 2015). In the case of TTL protein, TPR motifs are arranged in two tandem triplicates at the C terminus of the protein (Rosado et al., 2006). Another domain is the thioredoxin-like (TRXL) domain, which is homologous to the thioredoxin (TRX) domain. TRXL loses reductase activity due to a lack of a critical cysteine at the active site but is still considered to be important for protein stability (Ceserani et al., 2009; Amorim-Silva et al., 2019).

While searching for relevant interacting partners during a co-immunoprecipitation experiment, we encountered 421 proteins interacting, directly or indirectly, with TTL3 (Fig. S3, Table S3). In addition to enzymes and nucleic acid binding proteins involved in conventional cell activities, transporter proteins and cytoskeletal proteins are particularly prominent, which is consistent with the expression and localization of TTL3 in the cell (Fig. 6; Fig. 7). With the addition of 25 membrane-localized transferases, a total of 60 TTL3 interaction candidates participate in intracellular transport (Figure S3), five of which are clathrins that can construct coated vesicles to transport molecules within the cell and ten of which are Golgi associated proteins. The very abundant candidate interactors are related to the cytoskeleton, including 11 tubulins, 6 actins, and 6 cytoskeleton related proteins. According to Y2H and BiFC experiments, Tubulin 6 was the only confirmed stable and specific interaction. This physical interaction is indicated also by other datasets, where TTL3 was co-purified with tubulin as a candidate interactor (Hamada et al, 2013).

### TTL3 interacts with microtubules

TTLs are proteins without any target peptides directing them to cellular compartments (Amorim-Silva et al., 2019). Interestingly, the SUBA database (https://suba.live/factsheet.html?id=AT2G42580) predicts their possible localization in the nucleus and cytoskeleton (Hooper et al., 2017). The cytosolic localization is in accordance with our observation and other experimental data (Amorim-Silva et al., 2019). However, cellular localisation of transiently expressed *35S::TTL3-GFP* in tobacco leaves revealed clear microtubule associated pattern which was disrupted after microtubule destabilization by oryzalin (Fig. 6A-6C). An association with microtubules was also recorded in the transition domain of the RAM of stable Arabidopsis transformants. However, their visibility varies among independent transformant lines and might be obscured by a high cytoplasmic signal caused by fragments connected to GFP or free GFP in some of those (Fig. 6D; Fig. S4). Cortical microtubules decorated by TTL3 were found in *pTTL3::TTL3-GFP* transformed Arabidopsis plants in the elongation zone of the primary root (Fig. 6D; Fig. 7E - Fig. 7G), and after treatment with oryzalin, microtubule signals disappeared (Fig. 6E). There is also a conspicuous enrichment of TTL3-GFP protein in the phragmoplast of dividing cells. TTL3 distribution along individual microtubules is not homogenous and bead like structures were commonly observed moving along the microtubules (Fig. 7H; Video S4). The functional connection of known facts, namely, the relationship to BR signaling and close physical association with microtubule components of the cytoskeleton, opens up a rather wide set of hypotheses. For example cell wall material and cell wall remodelling related transport would fit also with promoter activity of the TTL genes and publicly available co-expression data.

The cytoskeleton and the intercellular transport it supports is critical for plant cell development and differentiation, and for plants to response and adaptation to biotic and abiotic stressors (Chen et al., 2008; Lozano-Durán et al., 2014; Garcia de la Garma et al., 2015). This is intriguing in connection to cell wall synthesis and its patterning. Vesicle traffic via the cytoskeleton delivers enzyme complexes and polysaccharide precursors to the cell surface (Kim and Brandizzi, 2014). On the plasma membrane, the transverse orientation of cellulose microfibers synthesized by cellulose synthase complex (CESA) depends on the organization of cortical microtubules (Paredez et al. 2006). Expression of TTL1 and TTL4 is spatially related to the formation of schizogenous intercellular spaces and/or specific differentiation of the cell walls of guard cells. Presence of TTL3 in the phragmoplast during cell plate formation and cell wall biogenesis is in accord with such expectation.

The executive role of TTL genes in cell division is unknown. A recent study showed that the microtubule-associated protein CLASP maintains cell proliferation through the brassinosteroid signaling pathway. The CLASP tether sorts vesicles to microtubules, while brassinosteroids (BR) change the distribution of CLASP and reorganize the microtubules. These are related to the inhibition of meristem activity, but do not affect cell elongation (Ruan et al., 2018). Similarly, TTL3 interacts with microtubules and is involved in intracellular transport, which may affect the activity of meristematic cells through the BR signaling pathway. Although, in the *ttl3* single mutant, no meristem defect was found. Shorter RAM and fewer cells in it were shown in the *ttl1ttl3ttl4* triple mutant (Fig. 3C-3D). These indicate the possible redundancy of TTL genes and their role in the development of RAM.

The co-expression data further supports the possibility of TTL3 playing a role in cell wall development, where genes for cell wall synthesis and modification are abundant (https://atted.jp/gene_coexpression/?gene_id=818858; www.genevestigator.com). TTL3 localisation was highly dynamic in the cell. The bead-like structures could be related to intracellular transport, for example, cell wall material. A new study shows that vesicle trafficking mediates the distribution of pectin in the cell wall, which in turn is essential for root clock function and subsequent LR formation (Wachsman et al., 2020). Pectin is synthesized in the form of highly methyl-esterified polymer in the Golgi and is delivered to the apoplast, where it can be de-methylesterified. The piecewise de-esterified homogalacturonan chains can be cross-linked by calcium, causing cell walls to harden and/or promote adhesion between cells (Zhang et al., 1992; Toyooka et al., 2009; Kim et al., 2015), whereas randomly de-esterified homogalacturonan can be degraded by pectinase, resulting in the loosening of the wall and/or cell separation (Goldberg et al., 1996; Daher and Braybrook, 2015; Yang et al., 2018). Without a way to regulate cell wall stiffness, the pectin methyl esterase 3 mutants (*pme3*) show a reduced number of LRs (Micheli 2001; Wachsman et al., 2020). Interestingly, PME3 is among the candidates for TTL3 interaction (Table S3). It is currently believed that the transportation of pectin, hemicellulose, and cellulose synthase complexes to the plasma membrane (PM) utilizes the conventional ER-Golgi-trans-Golgi network (TGN)-PM traffic route (Sinclair et al., 2018). Both pectin and hemicellulose are synthesized by enzymes located in the Golgi apparatus, and therefore need to be transported to the plasma membrane (McFarlane et al., 2014). The special role of Golgi and TGN in plants is the biosynthesis and sorting of cell wall components, including biosynthetic enzymes, structural proteins and matrix polysaccharides, hemicellulose, and pectin (Kim and Brandizzi, 2016). Among the candidate interaction proteins of TTL3, 10 proteins are related to the Golgi and TGN (Table S3). Therefore, TTL3 may interact with the Golgi to affect cell wall characteristics.

Moreover, there are eight aquaporins among the TTL3 interacting candidates (Table S3). The growth of LRP is caused by the relaxation of the cell wall and the expansion of the cell driven by its turgor pressure. The turgor pressure is maintained by the inflow of water. Aquaporins are membrane channels that promote the movement of water across cell membranes (Vilches-Barro and Maizel, 2015). Auxin can fine-tune the hydraulic characteristics of cells in LRPs and their overlying tissues by regulating the spatial expression of PIP genes. Over-expression and/or mutation of aquaporins significantly delayed the emergence of LR (Swarup et al., 2008; Péret et al., 2012).

In conclusion, we demonstrated that TTL3 interacts with microtubules, and that modulation of its expression changes the sensitivity of LRP development to brassinosteroids. Multiple TTL mutants are affected in RAM activity. TTL3 provides a novel bridge between these crucial elements of plant development, which are still unexplored, not only in the context of LRP development. Hopefully it will stimulate further research on the role of vesicular transport, microtubules, brassinosteroid signaling in cell division and lateral root development.

## Acknowledgement

Funding of Ministry of Education Youth and Sport (project NPUI LO1417) is gratefully acknowledged. The authors declare no conflict of interest. The Mexican enhancer and gene trap collection was funded by Consejo Nacional de Ciencia y Tecnología (CONACyT) and the Howard Hughes Medical Institute. The work in J.G.D. laboratory is partially supported by CONACyT grant A1-S-9236 and by DGAPA-PAPIIT-UNAM (grant IN200818). The authors thank Selene Napsucialy-Mendivil for technical help at the initial stage of the project.

## Supplementary files

**Table S1. List of primers used in this study.**

**Table S2. Categories of TTL3 interaction partners separated by co-immunoprecipitation.**

**Table S3. The candidate interactive partners of TTL3**. Total proteins were extracted from 2-week-old Arabidopsis seedlings expressing *pTTL3::TTL3-GFP* and *35S-GFP* (as a subtracted control). Protein complexes with *TTL3-GFP* and free GFP were isolated and analyzed by mass spectrometry.

**Video S1. The TTL3-GFP signal in tobacco.** Transient transformation of *Nicotiniana benthamiana* was carried out by *Agrobacterium tumefaciens* containing *35S::TTL3-GFP* plasmid and viral p19 post-translational silencing suppressor. High affinity of TTL3-GFP to microtubules was recorded and bead-like structures were found to move along the microtubule network.

**Video S2. The TTL3-GFP signal in Arabidopsis root transition zone.** Stable transformants of Arabidopsis *pTTL3::TTL3-GFP* show the network pattern typical of cortical microtubules and cytoplasmic localization in the transition zone of primary root.

**Video S3. The TTL3-GFP signal in Arabidopsis root transition zone after oryzalin treatment.** The Arabidopsis (containing *pTTL3::TTL3-GFP*) were treated with 100μM oryzalin, the changes of microtubules were recorded 10 min latter.

**Video S4. The TTL3-GFP bodies in differentiation zone of Arabidopsis root**. The mobile bodies in the differentiation zone of Arabidopsis root (containing *pTTL3::TTL3-GFP*).

**Video S5. The signal of TTL3-GFP and MAP4-RFP in Arabidopsis root transition zone.** The *35S::MAP4-RFP* (Microtubule-associated protein 4) and *pTTL3::TTL3-GFP* demonstrate spatial association of TTL3 protein and microtubules. Recorded in transition zone of primary root.

**Video S6. The bodies with TTL3-GFP signal moving along the microtubules.** The TTL3-GFP bodies signal moving along cortical microtubules (*pTTL3::TTL3-GFP* and *35S::MAP4-RFP*) in the differentiation zone of Arabidopsis primary root.

## References

Amorim-Silva, V., García-Moreno, A., Castillo, A.G., Lakhssassi, N., Valle, A.E., Pérez-Sancho, J., et al. (2019). TTL proteins scaffold brassinosteroid signaling components at the plasma membrane to optimize signal transduction in Arabidopsis. Plant Cell. 31(8), 1807–1828.

Azpiroz, R., Wu, Y., LoCascio, J.C., Feldmann, K.A. (1998). An Arabidopsis Brassinosteroid-Dependent Mutant Is Blocked in Cell Elongation. Plant Cell. 10, 219–230.

Banda J., Bellande K., von Wangenheim D., Goh T., Guyomarc’h S., Laplaze L., Bennett M.J. (2019). Lateral Root Formation in Arabidopsis: A Well-Ordered LRexit Trends Plant Sci. 24:826–839.

Bao, F., Shen, J., Brady, S.R., Muday, G.K., Asami, T., Yang Z. (2004). Brassinosteroids interact with auxin to promote lateral root development in Arabidopsis. Plant Physiol. 134 (4), 1624–1631.

Baskin, T.I., Beemster, G.T.S., Judy-March, J.E., Marga F. (2004). Disorganization of cortical microtubules stimulates tangential expansion and reduces the uniformity of cellulose microfibril alignment among cells in the root of Arabidopsis. Plant Physiol. 135 (4), 2279–2290.

Bhuiyan, N.H., Friso, G., Poliakov, A., Ponnala, L., and van Wijk, K.J. (2015). MET1 is a thylakoid-associated TPR protein involved in photosystem II supercomplex formation and repair in Arabidopsis. Plant Cell 27, 262–285.

Blatch, G.L., and Lässle, M. (1999). The tetratricopeptide repeat: a structural motif mediating protein-protein interactions. Bio Essays 21, 932–939.

Brückner, A., Polge, C., Lentze, N., Auerbach, D. and Schlattner, U. (2009). Yeast two-hybrid, a powerful tool for systems biology. International Journal of Molecular Sciences, 10(6), 2763–2788.

Butaye, K.M., Goderis, I.J., Wouters, P.F., Pues, J.M., Delauré, S.L., Broekaert, W.F., et al. (2004). Stable high-level transgene expression in Arabidopsis thaliana using gene silencing mutants and matrix attachment regions. Plant J. 39, 440–449.

Caesar K., Elgass K., Chen Z., Huppenberger P., Witthoft J., Schleifenbaum F., Blatt M.R., Oecking C., Harter K. (2011). A fast brassinolide regulated response pathway in the plasma membrane of Arabidopsis thaliana. Plant J., 66:528–540.

Catterou, M., Dubois, F., Schaller, H., Aubanelle, L., Vilcot, B., Sangwan-Norreel, B.S., et al. (2001a). Brassinosteroids, microtubules and cell elongation in Arabidopsis thaliana. I. Molecular, cellular and physiological characterization of the Arabidopsis bul1 mutant, defective in the Δ7-sterol-C5-desaturation step leading to brassinosteroid biosynthesis. Planta. 212, 659–672.

Catterou, M., Dubois, F., Schaller, H., Aubanelle, L., Vilcot, B., Sangwan-Norreel, B.S., et al. (2001b). Brassinosteroids, microtubules and cell elongation in Arabidopsis thaliana. II. Effects of brassinosteroids on microtubules and cell elongation in the bul1 mutant. Planta. 212, 673–683.

Ceserani, T., Trofka, A., Gandotra, N., Nelson, T. (2009). VH1/BRL2 receptor like kinase interacts with vascular-specific adaptor proteins VIT and VIK to influence leaf venation. Plant J. 57, 1000 – 1014.

Chaiwanon, J., and Wang, Z. Y. (2015). Spatiotemporal brassinosteroid signaling and antagonism with auxin pattern stem cell dynamics in Arabidopsis roots. Curr. Biol. 25, 1031–1042.

Chen, Y.N., Slabaugh, E., Brandizzi, F. (2008). Membrane-tethered transcription factors in Arabidopsis thaliana: novel regulators in stress response and development. Curr. Opin. Plant Biol. 11, 695–701.

Cho, H., Ryu, H., Rho, S., Hill, K., Smith, S., Audenaert, D., et al. (2014). A secreted peptide acts on BIN2-mediated phosphorylation of ARFs to potentiate auxin response during lateral root development. Nat. Cell Biol. 16, 66–76.

Clough, S.J., Bent, A. (1998). Floral dip: a simplified method for Agrobacterium - mediated transformation of Arabidopsis thaliana. Plant J. 16, 735–743.

Daher, F.B., Braybrook, S.A. (2015). How to let go: Pectin and plant cell adhesion. Front. Plant Sci. 6, 1–8.

De-Jesús-García R., Rosas U., Dubrovsky J.G. (2020). The barrier function of plant roots: biological bases for selective uptake and avoidance of soil compounds. Functional Plant Biology, 47(5), 383–397.

Delay, C., Imin, N., Djordjevic, M. (2013). Regulation of Arabidopsis root development by small signaling peptides. Frontiers in Plant Science 4, 352.

Deng, Q., Wang, X., Zhang, D., Wang, X., Feng, C., Xu, S. (2017). BRS1 function in facilitating lateral root emergence in Arabidopsis. Int. J. Mol. Sci. 18, 1549–1560.

Dittmer, H. G. (1937). A quantitative study of the roots and root hairs of a winter rye plant (Secale cereale). Am. J. Bot. 24, 417–420.

Du, Y.J., and Scheres, B., (2018). Lateral root formation and the multiple roles of auxin. J. Exp. Bot. 69 (2), 155–167.

Estrada-Luna, AA., Huanca-Mamani, W., Acosta-García, G., León-Martínez, G., Becerra-Flora, A., Pérez-Ruíz, R., and J-Ph. Vielle-Calzada. (2002). Beyond promiscuity: manipulating sexuality and apomixis in flowering plants. In Vitro Cellular and Developmental Biology - Plant 38, 146–151.

McFarlane, H. E., Doring, A., and Persson, S. (2014). The cell biology of cellulose synthesis. Annu. Rev. Plant Biol. 65, 69–94.

Fukaki, H., Tameda, S., Masuda, H. and Tasaka, M. (2002). Lateral root formation is blocked by a gain-of-function mutation in the SOLITARY - ROOT/IAA14 gene of Arabidopsis. Plant J. 29, 153–168.

Garcia de la Garma, J., Fernandez-Garcia, N., Bardisi, E., Pallol, B., Asensio-Rubio, J.S., Bru, R., et al. (2015). New insights into plant salt acclimation: the roles of vesicle trafficking and reactive oxygen species signalling in mitochondria and the endomembrane system. New Phytol. 205, 216–239.

Goh, T., Joi, S., Mimura, T., and Fukaki, H. (2012). The establishment of asymmetry in Arabidopsis lateral root founder cells is regulated by LBD16/ASL18 and related LBD/ASL proteins. Development 139, 883–893.

Goldberg, R., Morvan, C., Jauneau, A., Jarvis, M.C. (1996). Methyl-esterification, de-esterification and gelation of pectins in the primary cell wall. Prog. Biotechnol. 14, 151–172.

Gutierrez, R., Lindeboom, J.J., Paredez, A.R., Emons, A.M.C., Ehrhardt, D.W. (2009). Arabidopsis cortical microtubules position cellulose synthase delivery to the plasma membrane and interact with cellulose synthase trafficking compartments. Nat. Cell Biol. 11, 797–806.

Hamada, T.; Nagasaki-Takeuchi, N.; Kato, T.; Fujiwara, M.; Sonobe, S.; Fukao, Y.; Hashimoto, T. (2013) Purification and Characterization of Novel Microtubule-Associated Proteins from Arabidopsis Cell Suspension Cultures. Plant Physiology, 163, 1804–1816.

He, J.X., Gendron, J.M., Sun, Y., Gampala, S.S., Gendron, N., Sun, C.Q., Wang, Z.Y. (2005). BZR1 is a transcriptional repressor with dual roles in brassinosteroid homeostasis and growth responses. Science 307, 1634–1638.

He, J.X., Gendron, J.M., Yang, Y., Li, J., Wang, Z.Y. (2002). The GSK3-like kinase BIN2 phosphorylates and destabilizes BZR1, a positive regulator of the brassinosteroid signaling pathway in Arabidopsis. Proc. Natl. Acad. Sci. 99, 10185–10190.

Hochholdinger, F. and Zimmermann, R. (2008). Conserved and diverse mechanisms in root development. Curr. Opin. Plant Biol. 11, 70–74.

Hooper CM, Castleden I, Tanz SK, Aryamanesh, and Millar, AH (2017) SUBA4: the interactive data analysis centre for Arabidopsis subcellular protein locations Nucleic Acids Res. Jan 4;45(D1):D1064–D1074.

Hu, Z., Xu, F., Guan, L., Qian, P., Liu, Y., Zhang, H., et al. (2014). The tetratricopeptide repeat-containing protein slow green1 is required for chloroplast development in Arabidopsis. J. Exp. Bot. 65, 1111–1123.

Ivanov VB, Dubrovsky J. (2013) Longitudinal zonation pattern in plant roots: conflicts and solutions. Trends in Plant Science 18: 237–243.

Jacobsen, S.E., Binkowski, K.A., and Olszewski, N.E. (1996). SPINDLY, a tetratricopeptide repeat protein involved in gibberellin signal transduction in Arabidopsis. Proc. Natl. Acad. Sci. 93, 9292–9296.

Kang, Y.H., Breda, A., Hardtke, C.S. (2017). Brassinosteroid signaling directs formative cell divisions and protophloem differentiation in Arabidopsis root meristems. Development 144, 272–280.

Karimi, M., Bleys, A., Vanderhaeghen, R., Hilson, P. (2007). Building blocks for plant gene assembly. Plant Physiol 145, 1183–1191.

Kerppola, T. K. (2008). Bimolecular fluorescence complementation (BiFC) analysis as a probe of protein interactions in living cells. Annu. Rev. Biophys. 37, 465–487.

Kim, H., Park, P. J., Hwang, H. J., Lee, S. Y., Oh, M. H., and Kim, S. G. (2006). Brassinosteroid signals control expression of the AXR3/IAA17 gene in the cross-talk point with auxin in root development. Biosci. Biotechnol. Biochem. 70, 768–773.

Kim, S.J., and Brandizzi, F. (2014). The plant secretory pathway: an essential factory for building the plant cell wall. Plant Cell Physiol. 55, 687–693.

Kim, S.J., and Brandizzi, F. (2016). The plant secretory pathway for the trafficking of cell wall polysaccharides and glycoproteins. Glycobiology. 26, 940–949.

Kim, S.J., Held, M.A., Zemelis, S., Wilkerson, C., and Brandizzi, F. (2015). CGR2 and CGR3 have critical overlapping roles in pectin methylesterification and plant growth in Arabidopsis thaliana. Plant J. 82, 208–220.

Kim, S.K., Chang, S.C., Lee, E.J., Chung, W.S., Kim, Y.S., Hwang, S., Lee, J.S. (2000). Involvement of brassinosteroids in the gravitropic response of primary root of maize. Plant Physiol. 123 (3), 997–1004.

Kim, T., Lee, S.M., Joo, S., Yun, H.S., Lee, Y., Kaufman, P.B., Kirakosyan, A., Kim, S., Nam, K.H., Lee, J.S., Chang, S.C., Kim, S. (2007). Elongation and gravitropic responses of Arabidopsis roots are regulated by brassinolide and IAA. Plant Cell Environ. 30, 679–689.

Kotogány, E., Dudits, D., Horváth, G.V., Ayaydin, F. (2010). A rapid and robust assay for detection of S-phase cell cycle progression in plant cells and tissues by using ethynyl deoxyuridine. Plant Methods 6: 5.

Lakhssassi, N., Doblas, V.G., Rosado, A., Valle, A.E.D., Pose, D., Jimenez, A.J., et al. (2012). The Arabidopsis tetracopeptide thioredoxin-like gene family is required for osmotic stress tolerance and male sporogenesis. Plant Physiol. 158, 1252–1266.

Landrein, B., and Hamant, O. (2013). How mechanical stress controls microtubule behavior and morphogenesis in plants: history, experiments and revisited theories. Plant J. 75, 324–338.

Li, L., Xu, J., Xu, Z.H., Xue, H.W. (2005) Brassinosteroids stimulate plant tropisms through modulation of polar auxin transport in Brassica and Arabidopsis. Plant Cell. 17 (10), 2738–2753.

Liu, Y.G., Whittier, R.F. (1995). Thermal asymmetric interlaced PCR: automatable amplification and sequencing of insert end fragments from P1 and YAC clones for chromosome walking. Genomics 25, 674–681.

Lloyd, C.W., Clayton, L., Dawson, P.J, Doonan, J.H., Hulme, J.S., Roberts, I.N., Wells, B. (1985). The cytoskeleton underlying side walls and cross walls in plants: molecules and macromolecular assemblies. J. Cell Sci. 1985, 143–155.

Lozano-Durán, R., Bourdais, G., He, S.Y., Robatzek, S. (2014). The bacterial effector HopM1 suppresses PAMP-triggered oxidative burst and stomatal immunity. New Phytol. 202, 259–269.

Lucas M., Godin C., Jay-Allemand C., Laplaze L. (2008). Auxin fluxes in the root apex co-regulate gravitropism and lateral root initiation. J. Exp. Bot. 59: 55–66.

Maharjan, P.M., Schulz, B., Choe, S. (2011). BIN2/DWF12 antagonistically transduces brassinosteroid and auxin signals in the roots of Arabidopsis. J Plant Biol. 54, 126–134.

Malamy, J.E., and Benfey, P.N. (1997). Down and out in Arabidopsis: the formation of lateral roots. Trends Plant Sci 2, 390–396.

Marc J., Granger C., Brincat J., Fisher D., Kao T., McCubbin A., Cyr R. (1998) A GFP-MAP4 reporter gene for visualizing cortical microtubule rearrangements in living epidermal cells. Plant Cell 10, 1927–1940.

Marhavý, P., Montesinos, J.C., Abuzeineh, A., Van Damme, D., Vermeer, J.E.M., Duclercq, J., Rakusová, H., Nováková, P., Friml, J., Geldner, N., et al. (2016). Targeted cell elimination reveals an auxin-guided biphasic mode of lateral root initiation. Genes & Development, 30, 471–483.

Micheli, F. (2001). Pectin methylesterases: Cell wall enzymes with important roles in plant physiology. Trends Plant Sci. 6, 414–419.

Murashige, T., Skoog, F. (1962). A revised medium for rapid growth and bioassays with plant tissue culture. J. Plant Physiol. 15, 473–479.

Nakamura, A., Nakajima, N., Goda, H., Shimada, Y., Hayashi, K., Nozaki, H., Asami, T., Yoshida, S., Fujioka, S. (2006). Arabidopsis Aux/IAA genes are involved in brassinosteroid - mediated growth responses in a manner dependent on organ type. Plant J. 45, 193–205.

Paredez, A.R., Somerville, C.R., Ehrhardt, D.W. (2006). Visualization of cellulose synthase demonstrates functional association with microtubules. Science. 9, 1491–1495.

Pavelescu I., Vilarrasa-Blasi J., Planas-Riverola A., González-García M.P., Caño-Delgado A.I., Ibañes M. (2018) A Sizer model for cell differentiation in Arabidopsis thaliana root growth. Molecular Systems Biology 14, e7687.

Péret B., Li G., Zhao J., Band L.R., Voß U., Postaire O., Luu D.T., Da Ines O., Casimiro I., Lucas M., et al. (2012) Auxin regulates aquaporin function to facilitate lateral root emergence. Nat. Cell Biol. 14, 991–998.

Qiu, W., Park, J.W., Scholthof, H.B. (2002). Tombusvirus P19-mediated suppression of virus-induced gene silencing is controlled by genetic and dosage features that influence pathogenicity. Mol. Plant Microbe Interact. 15, pp. 269–280.

Rana, S., Hardtke, C.S. (2020). Plant Biology: Brassinosteroids and the Intracellular Auxin Shuttle. Curr. Biol., 30(9), R407–R409.

Rosado, A., Schapire, A.L., Bressan, R.A., Harfouche, A.L., Hasegawa, P.M., Valpuesta, V., et al. (2006). The Arabidopsis tetratricopeptide repeat-containing protein TTL1 is required for osmotic stress responses and abscisic acid sensitivity. Plant Physiol. 142, 1113–1126.

Roycewicz, P.S. and Malamy, J.E. (2014) Cell wall properties play an important role in the emergence of lateral root primordia from the parent root. J. Exp. Bot. 65, 2057–2069.

Sasse, J., Simon S., Gübeli C., Liu G.W., Cheng X., Friml J., et al. (2015). Asymmetric localizations of the ABC transporter PaPDR1 trace paths of directional strigolactone transport. Curr. Biol. 25, 647–655.

Scholl, R.L., May, S.T., Ware, D.H. (2000). Seed and molecular resources for Arabidopsis. Plant Physiol. 124, 1477–1480.

Sinclair, R., Rosquete, M.R., and Drakakaki, G. (2018). Post-Golgi trafficking and transport of cell wall components. Front. Plant Sci. 9: 1784.

Soukup, A. (2014). Selected Simple Methods of Plant Cell Wall Histochemistry and Staining for Light Microscopy; Humana Press: Totowa, NJ, USA, pp. 25–40.

Stoeckle, D., Thellmann, M., Vermeer, J.E. (2018). Breakout-lateral root emergence in Arabidopsis thaliana. Curr Opin Plant Biol 41, 67–72.

Sun, L., Feraru, E., Feraru, M.I., Waidmann, S., Wang, W., Passaia, G., Wang, Z.Y., Wabnik, K., Kleine-Vehn, J. (2020). PIN-LIKES coordinate brassinosteroid signaling with nuclear auxin input in Arabidopsis thaliana. Curr. Biol., 30, 1579–1588.

Sundaresan, V., Springer, P., Volpe, T., Haward, S., Jones, J.D.G., Dean, C., Ma, H., Martienssen, R. (1995). Patterns of gene action in plant development revealed by enhancer trap and gene trap transposable elements. Genes Dev, 9, 1797–1810.

Swarup, K. et al. (2008) The auxin influx carrier LAX3 promotes lateral root emergence. Nat. Cell Biol. 10, 946–954.

Szekeres, M., Németh, K., Koncz-Kálmán, Z., Mathur, J., Kauschmann, A., Altmann, T., et al. (1996). Brassinosteroids Rescue the Deficiency of CYP90, a Cytochrome P450, Controlling Cell Elongation and De-etiolation in Arabidopsis. Cell. 85, 171–182.

Torres-Martinez HH, Rodriguez-Alonso G, Shishkova S, Dubrovsky JG (2019) Lateral Root Primordium Morphogenesis in Angiosperms Frontiers in plant science 10:206.

Toyooka, K., Goto, Y., Asatsuma, S., Koizumi, M., Mitsui, T., Matsuoka, K. (2009). A mobile secretory vesicle cluster involved in mass transport from the golgi to the plant cell exterior. Plant Cell. 21, 1212–1229.

Vilches-Barro, A. and Maizel, A. (2015). Talking through walls: mechanisms of lateral root emergence in Arabidopsis thaliana. Curr Opin Plant Biol 23, 31–38.

von Wangenheim D., Banda J., Schmitz A., Boland J., Bishopp A., Maizel A., et al. (2020). Early developmental plasticity of lateral roots in response to asymmetric water availability. Nat. Plants 673–77.

Wachsman, G., Zhang, J., Moreno-Risueno, M. A., Anderson, C. T., Benfey, P. N. (2020). Cell wall remodeling and vesicle trafficking mediate the root clock in Arabidopsis.

Wang, Z.Y., Nakano, T., Gendron, J., He, J., Chen, M., Vafeados, D., Yang, Y., Fujioka, S., Yoshida, S., Asami, T., Chory, J. (2002). Nuclear-localized BZR1 mediates brassinosteroid-induced growth and feedback suppression of brassinosteroid biosynthesis. Dev. Cell. 2, 505–513.

Wilmoth, J.C., Wang, S., Tiwari, S.B., Joshi, A.D., Hagen, G., Guilfoyle, T.J., Alonso, J.M., Ecker, J.R. and Reed, J.W. (2005). NPH4/ARF7 and ARF19 promote leaf expansion and auxin -induced lateral root formation. Plant J. 43: 118–130.

Wolf S., van der Does D., Ladwig F., Sticht C., Kolbeck A., Schurholz A.K., Augustin S., Keinath N., Rausch T., Greiner S. et al. (2014). A receptor-like protein mediates the response to pectin modification by activating brassinosteroid signaling. Proceedings of the National Academy of Sciences, USA 111: 15261–15266.

Xuan W, De Gernier H and Beeckman T (2020) The Dynamic Nature and Regulation of the Root Clock. Development 147: 1–11.

Yang, Y., Sage, T.L., Liu, Y., Ahmad, T.R., Marshall, W.F., Shiu, S.-H., et al. (2011). CLUMPED CHLOROPLASTS 1 is required for plastid separation in Arabidopsis. Proc. Natl. Acad. Sci. 108, 18530–18535.

Yang, Y., Yu, Y., Liang, Y., Anderson, C.T., Cao, J. (2018). A profusion of molecular scissors for pectins: classification, expression, and functions of plant polygalacturonases. Front. Plant Sci. 9, 1–16.

Yin, Y., Wang, Z.Y., Mora-Garcia, S., Li, J., Yoshida, S., Asami, T., Chory, J. (2002). BES1 accumulates in the nucleus in response to brassinosteroids to regulate gene expression and promote stem elongation. Cell. 109, 181–191.

Yoshimitsu, Y., Tanaka, K., Fukuda, W., Asami, T., Yoshida, S., Hayashi, K., et al. (2011). Transcription of DWARF4 plays a crucial role in auxin-regulated root elongation in addition to brassinosteroid homeostasis in Arabidopsis thaliana. PLoS ONE 6, e23851.

Zar, J.H., (2010). Biostatistical Analysis, Ed 5th Edition. Pearson Prentice-Hall, Upper Saddle River, NJ.

Zhang, G.F., Staehelin, L.A. (1992). Functional Compartmentation of the Golgi Apparatus of Plant Cells. Plant Physiol. 99, 1070–1083.

